# BRCA2 C-terminal clamp restructures RAD51 dimers to bind B-DNA for replication fork stability

**DOI:** 10.1101/2024.09.21.614229

**Authors:** Michael A. Longo, Syed Moiz Ahmed, Yue Chen, Chi-Lin Tsai, Sarita Namjoshi, Xiaoyan Wang, Rajika L. Perera, Andy Arvai, Miyoung Lee, Li Ren Kong, Wilfried Engl, Woei Shyuan, Ziqing Winston Zhao, Ashok R. Venkitaraman, John A. Tainer, Katharina Schlacher

**Affiliations:** Department of Molecular & Cellular Oncology, UT MD Anderson Cancer Center, Houston, TX, USA; Texas A&M College of Medicine, Houston Methodist Willowbrook Hospital, Houston, TX, USA; The Cancer Science Institute of Singapore, National University of Singapore, Singapore, Singapore; Department of Cancer Biology, UT MD Anderson Cancer Center, Houston, TX, USA; Poseidon Laboratory, Pasadena, CA, USA; The Department of Integrative Structural & Computational Biology, The Scripps Research Institute, La Jolla, CA, USA; Medical Research Council Cancer Unit, University of Cambridge, Hills Road, Cambridge, UK; Department of Chemistry and Centre for BioImaging Sciences, National University of Singapore, Singapore; Mechanobiology Institute, National University of Singapore, Singapore; Institute of Molecular & Cell Biology Agency for Science, Technology and Research (A∗STAR), Singapore, Singapore

## Abstract

Tumor suppressor protein BRCA2 acts with RAD51 in replication-fork protection (FP) and homology-directed DNA break repair (HDR). Critical for cancer etiology and therapy resistance, BRCA2 C-terminus was thought to stabilize RAD51-filaments after they assemble on single-stranded (ss)DNA. Here we determined the detailed crystal structure for BRCA2 C-terminal interaction-domain (TR2i) with ATP-bound RAD51 prior to DNA binding. In contrast to recombinogenic RAD51-filaments comprising extended ATP-bound RAD51 dimers, TR2i unexpectedly reshapes ATP-RAD51 into a unique dimer conformation accommodating double-stranded B-DNA binding unsuited for HDR initiation. Structural, biochemical, and molecular results with interface-guided mutations uncover TR2i’s FP mechanism. Proline-driven secondary-structure stabilizes residue triads and spans the RAD51 dimer engaging pivotal interactions of RAD51 M210 and BRCA2 S3291/P3292, the cyclin-dependent kinase (CDK) phosphorylation site that toggles between FP during S-phase and HDR in G2. TR2i evidently acts as an allosteric clamp switching RAD51 from ssDNA to double-stranded and B-DNA binding enforcing FP over HDR.

## INTRODUCTION

Mutations affecting the 3418 amino acid <u>Br</u>east <u>Ca</u>ncer susceptibility protein 2 (BRCA2) predispose to hormone-driven and blood cancers, with the most frequent one resulting in carboxy terminal protein truncations ^1–5^. BRCA2 tumor suppressor function studies have largely been focused on its role in maintaining genome stability, which is accomplished by either DNA double strand break (DSB) repair pathway via Homology Directed Repair (HDR) restricted to G2 cell cycle phase or by DNA replication stability via DNA fork protection (FP) during S-phase ^6,7^. Yet, FP and HDR are genetically separable ^6,8^, whereby BRCA2 carboxy terminal RAD51 interactions are essential for FP but not for HDR. As replication errors appear responsible for two-thirds of the mutations in human cancers ^9^, an improved understanding of the BRCA2 C-terminus action at forks has fundamental importance.

Without mediator proteins, symmetric RAD51 dimers as a fundamental building block self-associate on both single stranded (ssDNA) and double-stranded DNA (dsDNA) ^10–15^. Assembly on flexible ssDNA occurs more rapidly than on the rigid dsDNA, owing to RAD51 oligomerization into helical filaments ^16^. In the HDR-active, ATP-bound state, the nucleoprotein filament is extended on ssDNA, which is thought to increase exposure of DNA bases to enable base homology search ^17^. In contrast, dsDNA binding triggers ATP hydrolysis, which renders RAD51 in an ADP-bound “compressed” form promoting RAD51 dissociation from DNA ^18–22^. How the interplay between nucleotide-bound states of RAD51 nucleoprotein filament complexes impact their distinct functions in HDR and FP has been enigmatic.

BRCA2 physically interacts with RAD51 recombinase by two general mechanisms: first via the 8 conserved BRC repeats encoded by exon 11 in human BRCA2 ^23,24^, and second via a conserved carboxy terminal RAD51 recognition domain (TR2) encoded within exon 27 (Figure 1A, B), which lacks sequence homology to the BRC repeats ^25–28^. Together the 8 BRC repeats enable both nucleation (BRC1-4) and stabilization (BRC5-8) of RAD51 filament growth on ssDNA, while suppressing dsDNA binding ^29–35^. In contrast, TR2 is thought to stabilize oligomeric RAD51 already assembled onto ssDNA ^25–27^, whereby a 36 residue peptide is minimally required for this distinct interaction (TR2i ^26^, Figure 1A, B).

**Figure 1.**
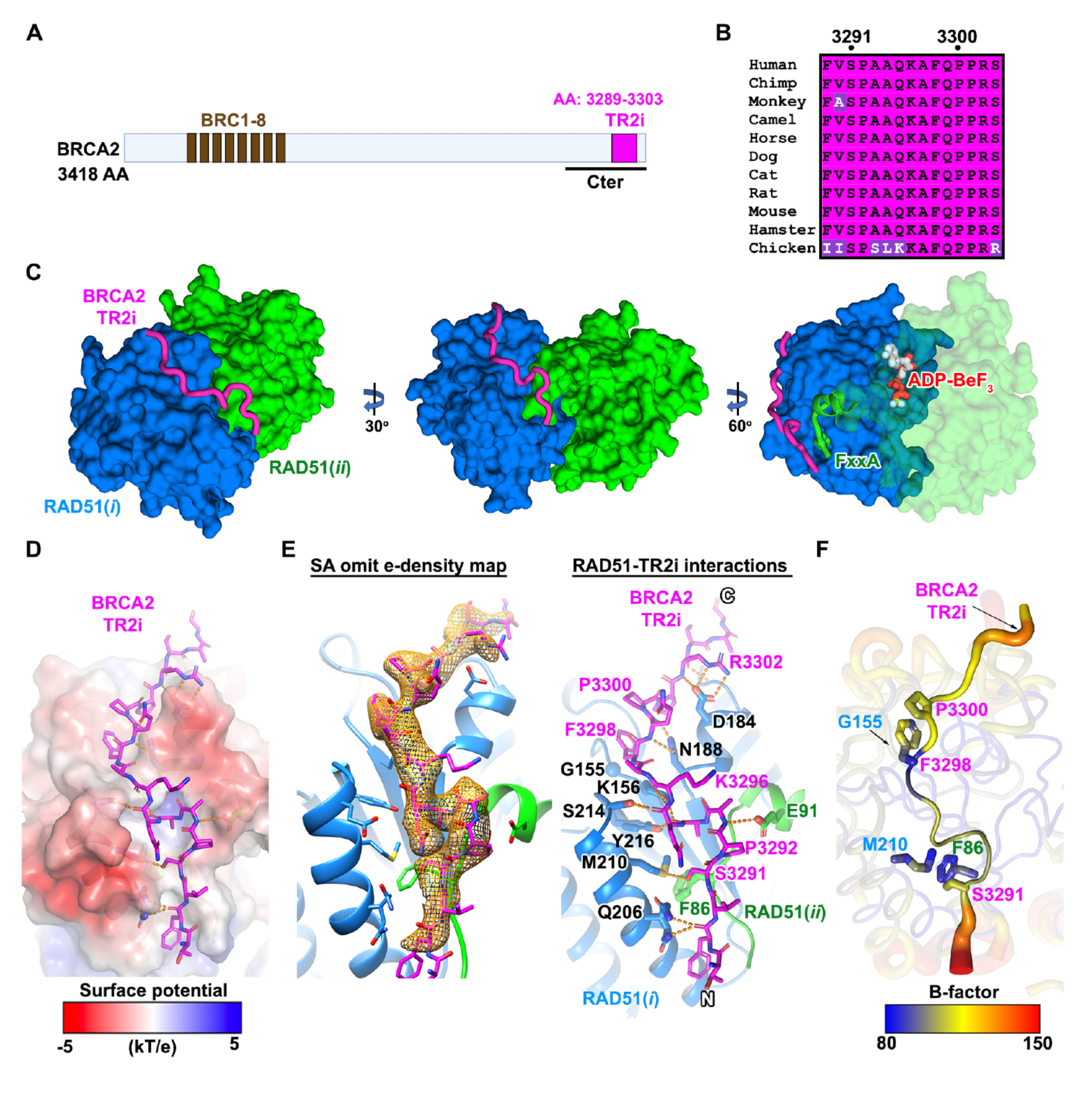
A conserved TR2i sequence and extended secondary structure binds within a groove across the RAD51 dimer and polymerization motif interface. (A) Full-length BRCA2 sequence schematic showing two separate RAD51 interaction regions: 8 N-terminal BRC motifs (brown, encoded by exon 11), and a non-homologous C-terminal RAD51 interaction region (magenta) originally mapped to amino acids (AA) 3270-3305 (TR2) and later refined to AA 3289-3303 as the minimally required interaction site (TR2i, magenta). **(B)** TR2i sequence alignment shows conservation for invariant (magenta) and variable (purple) sites. **(C)** The BRCA2 TR2i-RAD51 crystal structure (shown in three 30° rotations) reveals TR2i (residues 3289-3303, magenta) forms an extended curved clamping conformation within a groove across RAD51 oligomerization interface for RAD51(*i*) (blue) and RAD51*ii* (green) surface. In the rightmost image, TR2 caps the RAD51(*ii)* FxxA oligomerization motif (green Phe and ribbon) on a opposite face of the RAD51(*i*)-protomer from the ADP-BeF3 binding site at the RAD51 dimer interface (white and red sticks, center right). **(D)** RAD51 charged surface shows TR2i contacts are primarily by hydrophobic packing interactions and hydrogen bonds within a neutral groove, neighboring adjacent protruding acidic patches (red in the surface potential) that may facilitate onrate and orientation. **(E)** An objective BRCA2 TR2i Fo-Fc omit electron-density map (yellow mesh, left) shows defined main-chain conformation and side-chain positions (3σ contour level). A contact map depicts shape and packing complementarity plus detailed TR2i hydrogen bonds (dashes, right) within a groove across the RAD51 dimer. **(F)** Order and flexibility shown by TR2i crystallographic temperature factors (saturated tubes) overlying the RAD51 dimer (pale colors) reveal anchored ordered regions (blue/white narrow tube) centered at P3300, F3298 and at the S3291 tight turn. The marked flexibility at both ends of the TR2i (red widened tube) suggests that the TR2i-RAD51 dimer crystal structure contains the entire ordered interface.

RAD51-DNA filament stabilization is regulated by cyclin-dependent kinase (CDK), whereby phosphorylation of BRCA2 Serine at TR2 amino acids S3291/P3292 promotes filament disassembly ^26,27,36^. How this phosphorylation achieves RAD51 release from DNA has been unclear; yet, loss of exon 27, which harbors BRCA2-TR2i, predisposes to cancer, eliminates FP in cells, and renders cells mildly HDR-deficient ^37,38^. Strikingly, re-acquisition of BRCA2 C-terminus through frame shift mutation, which can fully restore FP and sub-optimally HDR^6,39^, confers cisplatin and PARP inhibitor resistance despite large deletions of other regions ^40,41^. In principle therapeutic resistance to PARP inhibitor may occur by restoration of FP, HDR, or both^42^. Mutation of CDK phosphorylation site S3291 to alanine abrogates the TR2i-RAD51 interaction and BRCA2 stabilizing effect on RAD51-DNA ^26,36^. Reintroducing S3291A mutant BRCA2 to cells with a C-terminally truncated BRCA2 rescues the mild HDR defect, but replication fork instability persists ^6^, supporting an exquisite essentiality of TR2i-RAD51 for FP.

A central unanswered question has therefore been how the unique TR2i-RAD51 interaction motif differentiates FP from HDR compared to known structural, biochemical and cellular BRC repeat functions. Here we define the TR2i-RAD51 X-ray crystal structure at 2.7 Å resolution (Table S1), which unveils a pivotal three-dimensional TR2i interface with the ATP-analogue bound RAD51 dimer. Our structural analysis was enabled by the atomic detail of this X-ray crystal structure in the absence of DNA, that is before it nucleates onto DNA. Combining the structural results with cellular and molecular mutation analyses, the data identifies BRCA2 TR2i as an allosteric clamp reshaping RAD51 dimer into an unprecedented conformation that changes RAD51 DNA substrate specificity to enforce ds and B-DNA binding. This finding defines its role in maintaining replication fork stability and separates its biophysical role for FP from HDR. The combined data thereby changes our understanding of how BRCA2 accomplishes two mutually exclusive RAD51 loading functions, which are controlled by a CDK phosphorylation switch that defines its role in maintaining replication fork stability.

## RESULTS

### BRCA2 TR2i binds across RAD51 dimer interface

RAD51 oligomerization is accomplished by critical interactions of the FxxA polymerization motif (PM) binding within a hydrophobic pocket of the C-terminal globular domain of an adjacent RAD51 protomer (Figure S1A). To study BRCA2-RAD51 interactions distinct to BRCA2 TR2i, we isolated an obligate RAD51 dimer incapable of continued RAD51 oligomerization (Figure S1A-D).

To achieve this, we truncated one RAD51 protomer subunit (RAD51(*i*), blue) beyond the FxxA polymerization motif (PM), which prevents forward N-terminal oligomerization, while retaining the globular RAD51 dimer interaction domains enabling C-terminal RAD51 binding (Figure S1B). We combined this with a second protomer subunit (RAD51(*ii*)), which retains both the FxxA polymerization and the globular domain but added a C-terminal polymerization blocking motif to prevent potential Rad51 oligomerization in the reverse (C-terminal) direction (Figure S1B). This design generates a defined RAD51(*i*)-RAD51(*ii*) dimer nucleation unit with a fully intact ATP-binding site and consecutive DNA binding loops. Given the negligible role of RAD51 N-terminal domain in interaction with BRCA2 C-terminus ^26,27^, we truncated the N-terminus of the protomers to aid crystallization of the protein complexes. This obligate RAD51 dimer produced a stable complex with a conserved BRCA2 region minimally required for RAD51 interactions with BRCA2 C-terminal (TR2i, amino acids 3289-3303 ^26^).

By comparative computational modeling this constitutive RAD51 dimer was designed to allow for known conformations and nucleotide-bound states within a RAD51:RAD51 interface. We therefore grew crystals of BRCA2 TR2i-RAD51 dimer in complex with the ATP analog ADP-BeF_3._ These crystals diffracted to a 2.7 Å resolution, whereby refined electron density maps allowed placement and definition of the TR2i-RAD51 dimer structure in atomic detail (Figure 1C and Figures S1D and S1E). Of note, the TR2i-RAD51 dimer structure formed crystal complexes without DNA despite the presence of poly-dT in the crystallization solution, supporting the strength of the peptide-protein interaction that occurs prior to DNA binding. The refined coordinates show one TR2i-RAD51 dimer within the asymmetric unit. The two ATP sites each bind the ATP analog ADP-BeF_3_ and are superimposable (Figure S1E), confirming ATP-binding site conservation. The experimental crystal structure furthermore validates the TR2i sequence as the entire interface region, as the residue ends become disordered as they reach the edges of the RAD51 subunits, consistent with the overall disorder of exon27 ^43^.

The TR2i-RAD51 dimer structure shows that the conserved BRCA TR2i sequence 3289-3303 forms an extended secondary structure with complementary interactions spanning across a groove in the RAD51 ATPase dimer interface (Figures 1C-E). TR2i directly overlays the FxxA polymerization motif ^44^ that buries the hydrophobic pocket of the adjacent RAD51 ATPase domain (Figures 1C and 1E). A recent publication on a cryo-EM structure with low resolution at the TR2-RAD51 interface suggested the importance of the RAD51 acidic patch interactions with TR2^45^. The atomic detail revealed by the crystal structures shows that the solvent-exposed charge interface has limited charge complementarity (Figures 1D). Instead, hydrophilic TR2i residues primarily point away from the interface, providing flexibility that enables allosteric action. In contrast, hydrophobic residues and main-chain atoms form complementary interactions within the interface groove (Figure 1D), providing stability for the interaction within the groove sheltered from bulk solvent. The objective electron density map with the TR2i omitted (Figure 1E, left) verifies that all interactions identified are defined by the experimental structure data (Figure 1E, right). The temperature factor map reveals flexibility in the TR2 edge (Figure 1F, red), suggesting the solvent-exposed charged interaction at the outer regions is secondary. In contrast, the main chain and hydrophobic side chains are highly ordered (Figure 1F, yellow and blue), confirming that these interactions, rather than those with the charged patch, predominate in forming the TR2-RAD51 interface.

### TR2i restructures the RAD51 dimer enabling B-form DNA binding

To better understand the effect of TR2i on the ATP-bound RAD51 dimer, we compared the structure with the published structure of the biologically active ATP-bound RAD51 dimer^46,47^ (Figures 2A and 2B,). Strikingly, TR2i restructures the RAD51 dimer into a conformation not previously reported for RAD51 (Figure 2, Movies S1-S3).

**Figure 2.**
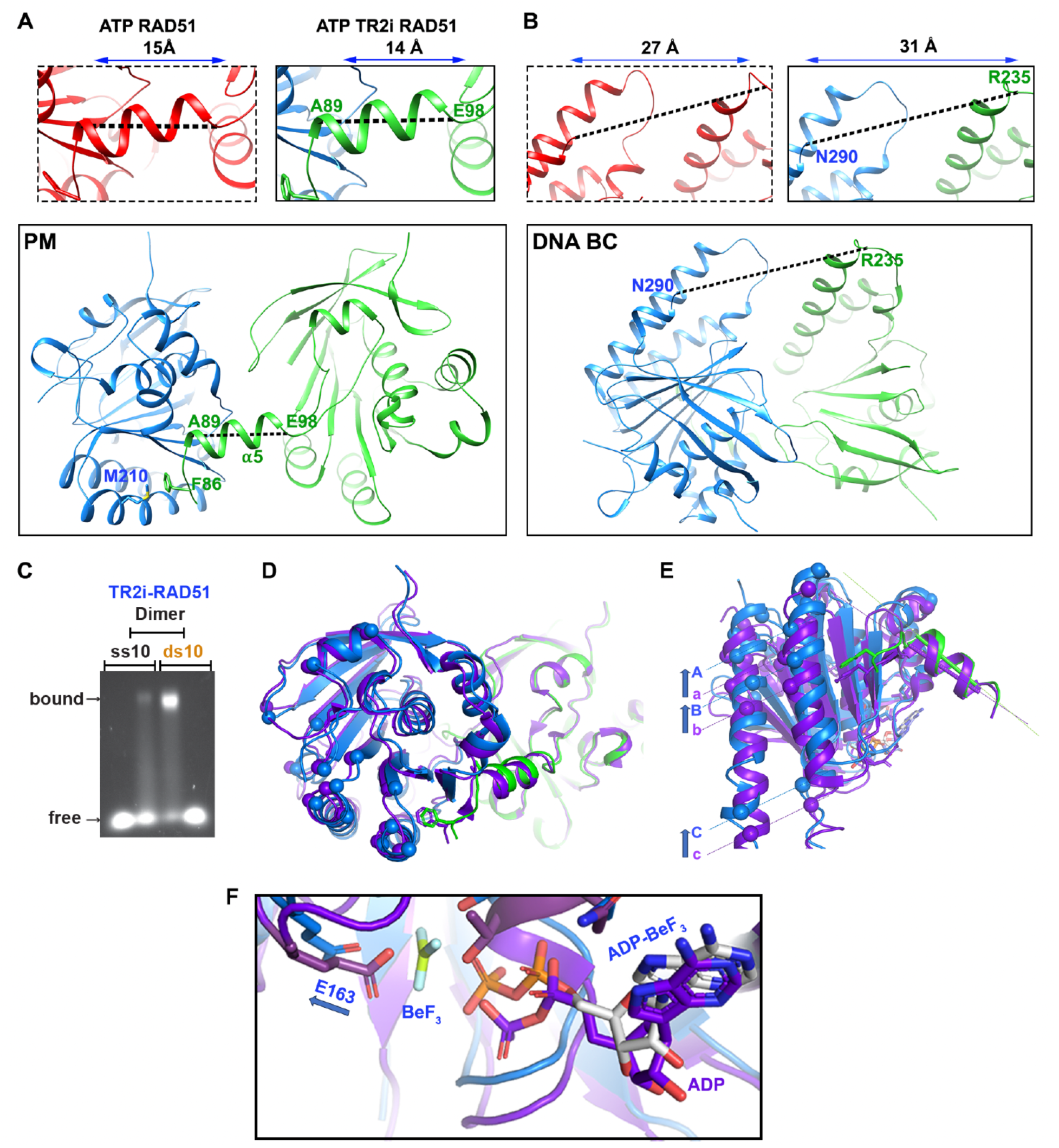
BRCA2 TR2i binding restructures the RAD51 dimer. **(A)** Top, RAD51(*ii*) PM *α*-helix is compressed (14Å green, right) compared to the ATP-bound RAD51 dimer structure without TR2i (15 Å red, left, PDB: 5NWL chains E,F, top). Bottom, TR2i-bound RAD51 dimer view (TR2i not shown) depicting the green helix connection to the M210 (blue) –F86 (green) polymerization motif (PM) interaction. **(B)** DNA binding channel (DNA BC) between RAD51 protomer subunits expands for TR2i bound to RAD51 dimer (31Å between RAD51(*i*) and (*ii*), top right) compared to that of the ATP-bound RAD51 dimer structure without TR2i (27Å, red, top left). Bottom, TR2i-bound RAD51 dimer (TR2i not shown) showing distance measured for Loop 1 RAD51(*ii*) to labeled RAD51(*i*) Loop 2 N290 that form the base of the DNA binding groove. **I** Representative electrophoretic mobility shift assays (EMSA, from > 3 experiments) shows that TR2i-RAD51 dimer binds more avidly to double-stranded DNA (ds, 10 bp, right) compared to single-stranded DNA (ss, 10 bp, left). **(D)** Structural alignment of the TR2i-RAD51 dimer with previously reported ADP-bound, compressed dimer (purple, PDB: 8BSC). RAD51(*i*) protomer (blue) appears to align in space and orientation with an ADP-compressed subunit (purple) from a top vieI**(E)** Rotating TR2i-RAD51 backward on a horizontal axis by 90° from the **D** orientation shows that the RAD51(*i*) protomer (blue) is uniquely shifted upwards (blue arrows) in the plane of the page for the major helices and the associated RAD51(**i*i*) PM motif (green) relative to the ADP-bound, compressed dimer (purple). **(F)** Close-up of ATP binding sites shows that the TR2i-RAD51i can accommodate the gamma-phosphate position (BeF_3_, blue) of the ATP analogue (“ADP-BeF_3_”, white carbon blue sticks) with E163 (blue) being shifted compared to the E163 in the ADP-bound RAD51 structure (purple and purple stick).

α-helix 5 of the RAD51(ii) protomer, which is part of the PM that acts as an interlocking handle between the RAD51 protomers, slightly compresses in the presence of TR2i (Figure 2A, 15 to 14 Å). As the PM interface compresses, the regions on the opposite side of the RAD51 dimer widen, a site that contains the DNA binding channel (Figure 2B). Thus, TR2i binding to ATP-RAD51 allosterically clamps the RAD51 dimer on one side while widening the DNA binding channel on the opposite side.

The TR2i-induced opening of the DNA channel suggests that the reshaped RAD51 dimer may exhibit a binding preference for more rigid and base-stacked B-form DNA. Strikingly, we observe TR2i-RAD51 dimers bind dsDNA much more readily than ssDNA (Figure 2C, 71% dsDNA compared to 12% ssDNA). Additionally, TR2i enhances RAD51 DNA binding to hyper-short ssDNA (Figures S2A and 2B). Taken together, the data shows BRCA2 TR2i-RAD51 dimer changes DNA substrate specificity of the RAD51 dimer to favor dsDNA and hyper-short B-DNA binding.

Double-stranded (ds) DNA binding is classically associated with ADP-RAD51 filament states after ATP hydrolysis, which promotes RAD51 compression, inactivation, and disassembly from DNA. While the RAD51(*i*) protomer in a top-view appears to align well with the previously reported structure of the compressed ADP-RAD51 protomer (Figure 2D), a 90-degree rotation reveals that the RAD51 protomer in the presence of TR2i is shifted (Figure 2E, blue) compared to ADP-RAD51 (Figure 2E, purple). Importantly, this unique TR2-RAD51 dimer conformation accommodates the γ-phosphate of the ATP, which otherwise clashes with the Glutamate 163 as seen in the ADP-RAD51 structure (Figure 2F). Thus, TR2i promotes a unique RAD51 conformation that allows dsDNA binding in the presence of ATP, so preventing disassembly from dsDNA as otherwise observed with ADP-bound RAD51.

### Main-chain interactions reveal two interface triads between BRCA2 TR2 and RAD51 dimer

The highly ordered main chain forms two proline driven triads defining TR2-RAD51 dimer interactions (Figure 3). TR2-RAD51 is controlled by CDK phosphorylation at BRCA2 S3291^36^. The structure reveals that S3291 also is critical to the BRCA2 TR2-RAD51 interaction as part of a (BRCA2 S3291: RAD51(*i*) M210 : RAD51(*ii*) F86) triad (Figure 3A and 3B), explaining how phosphorylation can act as a molecular switch. This triad positions the S3291 hydroxyl to hydrogen bond to M210 sulfur (Figures 3A and 3B). In turn, the RAD51(*i*) M210 side chain caps its PM binding pocket and so secures the F86 of the adjacent RAD51(*ii*) within the pocket (Figure 3B). M210 is conserved among RAD51 homologs across species, but unique to RAD51 compared to the human RAD51 paralogs (Figure S3B), supporting its distinct function in human RAD51-BRCA2 TR2i interaction.

**Figure 3.**
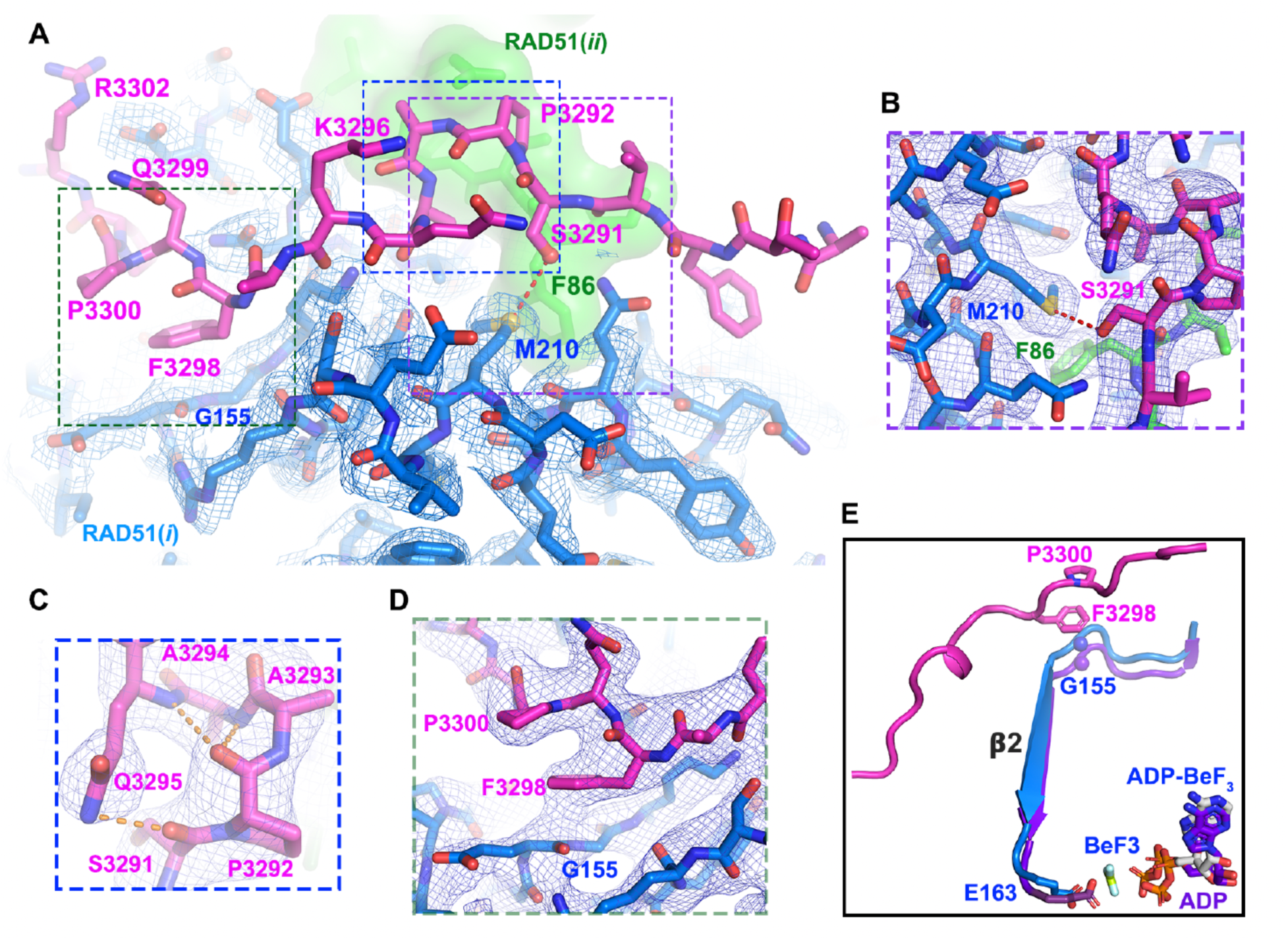
Ordered residue triads link TR2i binding to the RAD51 PM motif and ATP site to anchor the TR2i-RAD51 complex to RAD51 dimer conformation. Electron density maps are shown (2Fo-Fc 1σ contours, blue mesh). **(A)** Overview density map for RAD51(*i*) with TR2i-RAD51 dimer interaction triads. RAD51(*i*) (blue carbon sticks) and RAD51(ii) (green carbon sticks) show interactions with TR2i (magenta carbon sticks) with dashed boxes identifying the P3292 tight turn (top center) and the two critical triad anchor interactions: the P3300:F3298:G155 triad (left) and S3291:M210:F86 triad (right). The P3292-A-A-Q tight turn directs S3291 (red oxygen, purple carbon sticks) to hydrogen bond (red-dash) with RAD51(*i*) M210 (blue carbon sticks, yellow sulfur) stabilizing RAD51(*ii*) F86 (green sticks) in the PM motif via the S3291:M210:F86 triad (center right). BRCA2 F3298 stacks between P3300 (magenta carbon sticks, center left) and RAD51(*i*) G155 amide backbone (blue sticks) stabilizing the complex. **(B)** S3291:M210:F86 triad closeup. S3291 anchors TR2i N-terminal end via the triad interactions. S3291 (red oxygen, magenta carbon sticks) hydrogen bonds (red-dashes) to RAD51(*i*) M210 (blue carbon sticks, yellow sulfur) further stabilizing the underlying RAD51(*ii*) F86 (green carbon sticks) as the linchpin in the PM interface. **(C)** P3292-A-A-Q tight turn (yellow dash hydrogen bonds) closeup showing TR2i conformation directs S3291 side chain downward toward RAD51(*i*) M210 thioester. **(D)** P3300:F3298:G155 triad closeup. F3298 stacks between P3300 and amide backbone of RAD51(*i*) G155 (blue sticks) via π-π interactions anchoring the complex to RAD51(*i*) conformation. **(E)** P3300:F3298:G155 triad is positioned to network TR2i binding through RAD51(*i*) ß2 strand and E163 with the ATP-gamma phosphate.

At the RAD51(*i*) M210:TR2i interface, BRCA2 TR2i forms a tight turn initiated with S3291 (Figure 3C). The P3292 carbonyl oxygen hydrogen bonds to the amine of Q3295, the final residue of the turn, as is characteristic of tight turn structures ^48^. Further stabilizing this turn, the P3292 carbonyl group also hydrogen-bonds with other backbone amine groups within the turn (Figure 3C), consistent with the role of proline as instigator and stabilizer of peptide turns ^48–50^.

TR2i residue F3298, adjacent to the tight turn, forms the start of the second TR2-RAD51 interaction triad (BRCA2 F3298 : BRCA2 P3300 : RAD51 G155). Consistent with F3298 being necessary for TR2i-RAD51 dimer interaction^43^, it stacks on the backbone peptide bond of RAD51(*i*) G155 via a pi-pi interaction (Figure 3D), where G155 is conserved only across RAD51 homologs but not RAD51 paralogs (Figure S3A). P3300 sandwiches the F3298 with a predicted C-H/χ interaction ^51,52^. Notably, the P3300:F3298:G155 triad is positioned to network TR2i binding through RAD51(*i*) ß2 strand and E163, suggesting importance of this triad for the unique conformation with allosteric accommodation of the ATP-γ-phosphate analog (Fig. 3E).

Together, the C-terminal anchor F3298 and N-terminal anchor S3291 for the tight turn form the highly ordered region of the TR2i-RAD51 interface (Figure 1F and 3A). Outside these TR2i triad anchor points, the temperature-factors of the interface increase, indicating higher flexibility ^53^. So these TR2i structural elements define a bipartite RAD51 dimer interface for the BRCA2 C-terminus.

### S3291:M210:F86 and F3298:P3300:G155 triads are critical for BRCA2 TR2i-RAD51 protein interaction

The crystals without bound DNA and the structure obtained with the TR2i-RAD51 dimer suggest a strong protein-peptide interaction. Consistently, wild-type (WT) RAD51 co-elutes with His-tagged TR2 when purified by nickel chromatography, confirming a robust interaction between them (Figure 4A). Following these results, we devised a sequential chromatography purification method to test the structure implications for BRCA2 TR2-RAD51 and RAD51 dimer interactions further experimentally.

**Figure 4.**
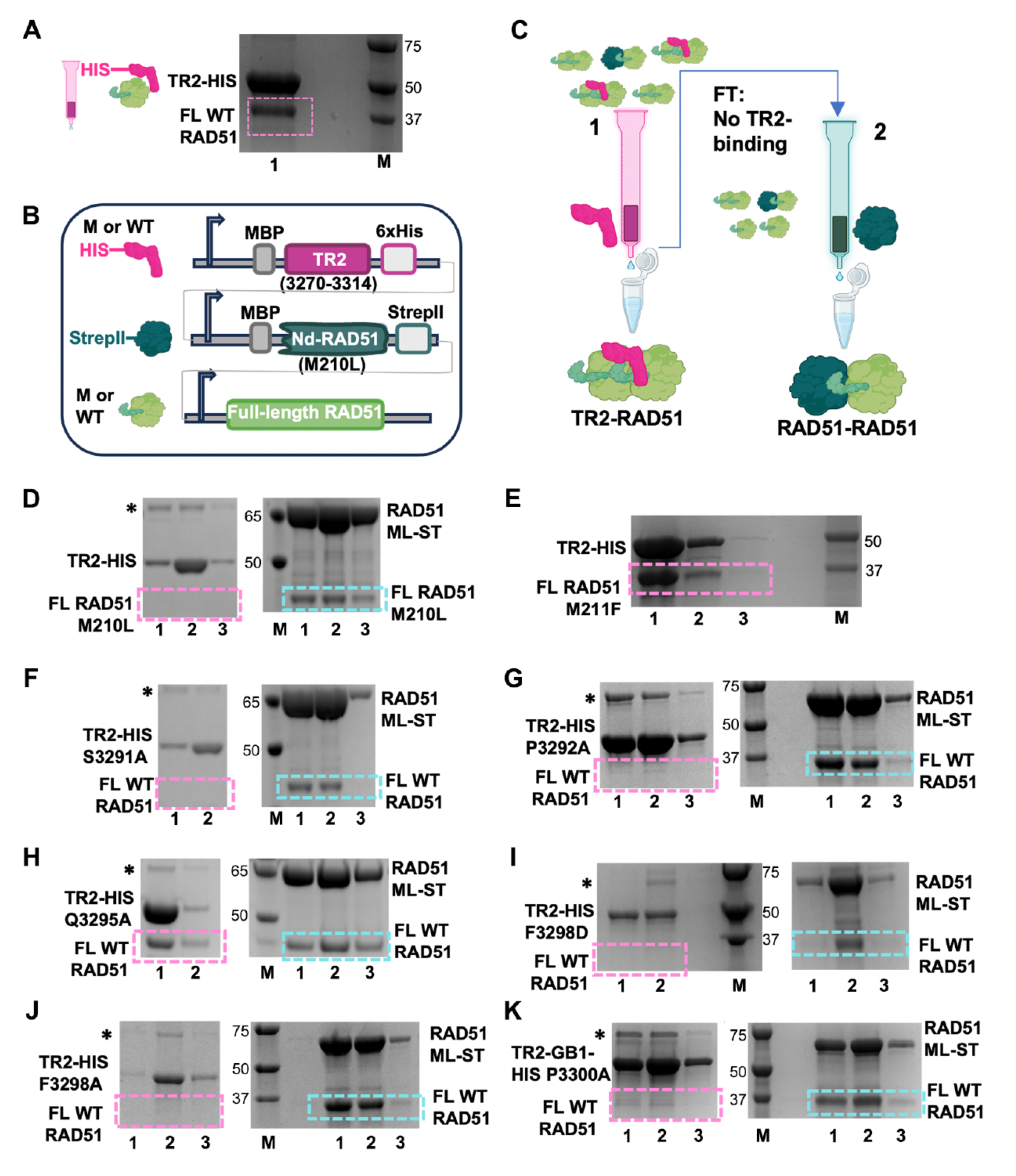
Biochemical analyses support the pivotal importance of the interface residue triads to BRCA2 TR2i-RAD51 complex stability. **(A)** Coomassie protein gel of eluents (numerical label 1) from Nickel 2+ chromatography column after running a cell lysate of His-tagged TR2 co-expressed with full length (FL) wild-type (WT) RAD51. M, protein size marker. **(B)** Expression construct used in 2-step purification method. His, histidine tag; Nd, N-terminal truncated; MBP, Maltose binding protein; M, Mutant; WT, wild type. **(C)** Schematic for 2-step purification method. FT, flowthrough. **(D)** Coomassie protein gel of eluents (numerical labels 1-3) from Nickel 2+ chromatography column (expected RAD51 gel band in magenta dashed box) and from the Strep-Tactin Sepharose column (expected RAD51 gel band in blue dashed box) of lysates from expression constructs containing WT TR2 and full-length RAD51 M210L. **(E)** WT TR2 and full-length RAD51 M211F. **(F-K)** Full-length wild-type RAD51 with mutant TR2 S3291A (**F**), P3292A (**G**), Q3295A (**H**), F3298D (**I**), F3298A (**J**), or P3300A (**K**). The TR2 P3300A expression construct additionally contains a B1 domain of streptococcal protein G (GB1) to prevent degradation of this unstable non-RAD51 interacting peptide.

For this we co-expressed His-tagged TR2 with full-length RAD51 together with an N-terminally truncated StrepII-tagged RAD51 M210L (Figure 4B). In a first purification step, His-TR2 plus any RAD51 bound to TR2 will elute together from the nickel chromatography column. Thus, introducing mutations in either TR2 or full length RAD51 tests the importance of a given residue to the TR2-RAD51 interactions by following their ability to co-elute (Figure 4C, 1). In a second step serving as an internal control, the flow through containing both full-length RAD51 and StrepII-tagged RAD51 M210L unable to bind to TR2 is applied to a StrepTactin column. Eluants of this column contain the StrepII-tagged RAD51 and any full-length RAD51 that retains interactions with the Strep-II tagged protein. The StrepII tagged RAD51 construct is N-terminally truncated to prevent forward polymerization and so avoid multi-valent interactions. Thus, co-eluants from this column test the C-terminal RAD51-RAD51 interactions (Figure 4C, 2). This system both tests the effect of a residue change on RAD51-RAD51 interactions when introducing mutations in full-length RAD51 and serves as an internal control for the nickel column results.

By using this co-expression and purification scheme, we tested the importance of the S3291:M210:F86 triad by introducing a M210L mutation into both RAD51 constructs. Despite leucine being the least disruptive mutation, RAD51 M210L does not co-elute with His-TR2i (Figure 4D, pink dashed line). Yet, it is proficient in binding to RAD51 to form dimers, as it co-elutes with StrepII-RAD51 (Figure 4D, blue dashed line). In contrast, mutation of the adjacent residue M211 had no effect on the TR2-RAD51 interaction (Figure 4E), validating M210 as critical to TR2i-RAD51 interaction. Further confirming structural observations, WT RAD51 is unable to co-elute with either TR2 S3291A (Figure 4F) or with TR2 P3292A (Figure 4G). Yet, mutation of Q3295A affecting the tight turn (Figure 3C) is not disruptive, as the TR2-RAD51 interaction remains intact (Figure 4H). To test the second triad, we introduced mutations in TR2 at F3298 and P3300. Similar to the S3291:M210:F86 triad mutations, TR2 F3298A, F3298D or P3300A all disrupt the TR2i-RAD51 interaction (Figures 4I, 4J, 4K), supporting that the F3298:P3300:G155 pi-stacking triad too is a critical second site defining a bipartite TR2i-RAD51 interface. Together these data confirm the specificity and importance of the bipartite S3291:M210:F86 and F3298:P3300:G155 triads for the TR2-RAD51 interactions.

### TR2i-RAD51 interface mutations abolish fork protection without affecting HDR

To test the role of the structurally defined interface for the BRCA2 TR2i-RAD51 interaction in the context of the cell, we introduced homozygous point mutations in the BRCA2 or RAD51 loci (BRCA2 P3292L, RAD51 M210L or RAD51 M210I, respectively) in the HeLa Kyoto cell line by CRISPR-Cas9 engineering confirmed by Sanger sequencing (Figure S4). The RAD51 M210I mutation is a cancer-associated variant with unreported clinical significance in ClinVar ^54^, and the BRCA2 P3292L mutation is classified as a ‘variant of unknown significance (VUS)’ or “benign” ^55–57^. As controls for HDR deficiency, we used BRCA2^Tr/Tr^ cells, which carry mutations within exon 11 leading to premature translation termination and deletion of the BRC repeats as well as TR2 and DNA binding motifs^58^.

BRCA2 C-terminal RAD51 interaction is essential for FP^6^. Consistent with the RAD51 fork-stability results, all BRCA2 TR2i-RAD51 interface mutations were significantly defective in FP as seen by the degradation of nascent DNA tracts following HU-mediated stalling observed by the DNA fiber assay (Figure 4A).

The combined data suggest severe defects of the point mutations in TR2-RAD51 binding. To test if these point mutations affect HDR-mediated DSB repair, we next employed the CRISPR-Cas9/mClover-LMNA1 assay. As expected, the BRCA2^Tr/Tr^ cells were severely deficient in HDR (Figures 5B-5D), consistent with the sensitivity observed when treating these cells with the PARP inhibitor Olaparib (Fig. S5A). In stark contrast, BRCA2 TR2i P3292L, RAD51 M210L and RAD51 M210I mutations showed no impacts on HDR with mutants showing efficiencies similar to wild-type cells (Figures 5B-5D). Furthermore, HeLa Kyoto cells carrying the aforementioned point-mutations in either BRCA2 or RAD51 survived Olaparib challenge comparably to WT BRCA2/RAD51 cells (Figure S5A.). Taken together, the data show that residues important for stable BRCA2 TR2i-RAD51 interactions are dispensable for RAD51 reactions during HDR but crucial for FP during DNA replication fork stalling.

**Figure 5.**
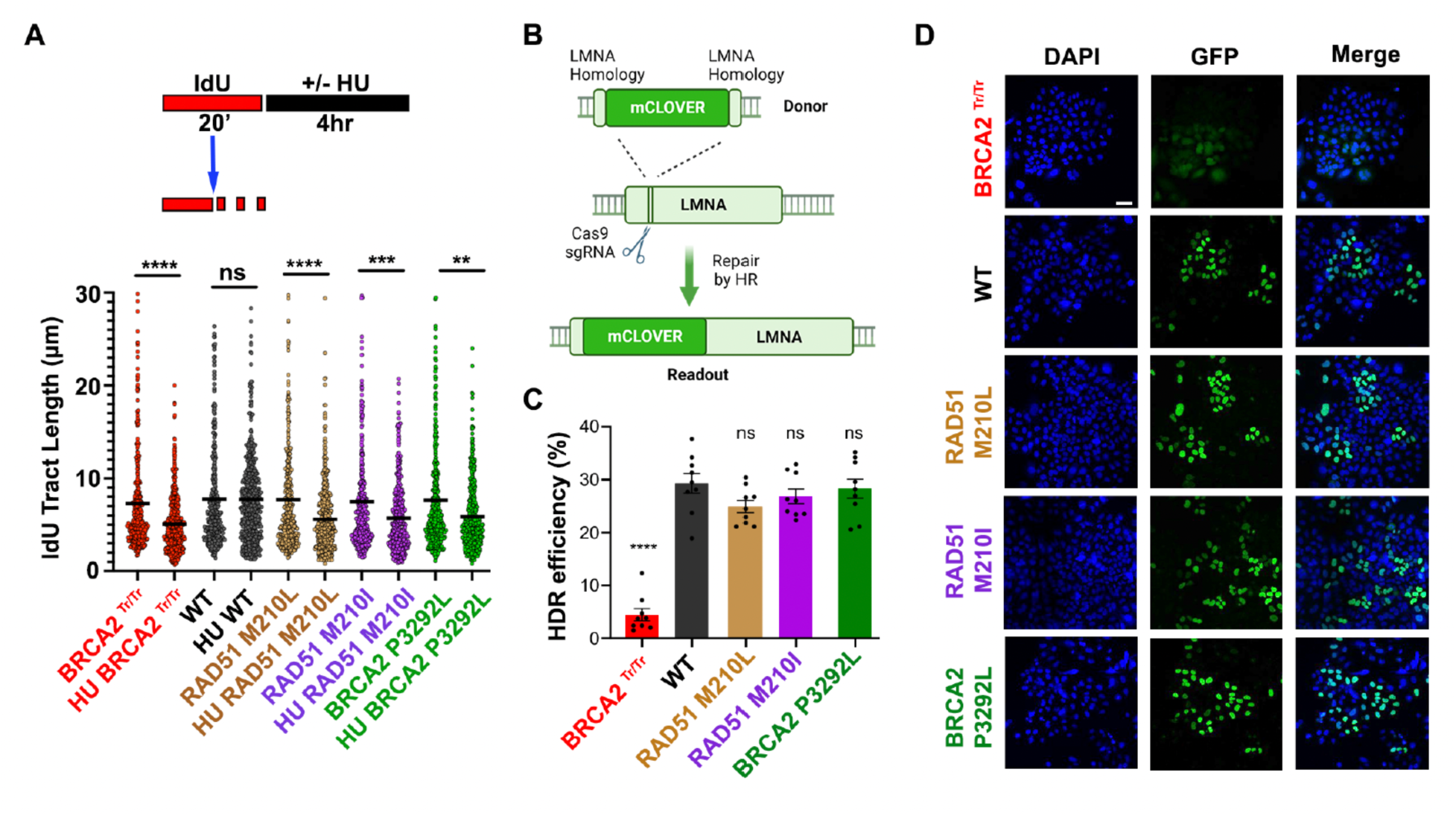
BRCA2 and RAD51 mutations impacting the TR2i-RAD51 dimer interaction cause FP defects despite HDR proficiency. **(A)** FP analyses in WT, RAD51 mutant, and BRCA2 mutant cells. Single-molecule DNA fiber analysis of nascent IdU tract length size distribution in WT and mutant HeLa Kyoto cells with and without HU treatment as a measure of fork protection (n>400, two independent biological experiments). Bar represents median. Two-tailed Mann-Whitney t test was used to determine *P* values for statistical analysis. \*\**P*< 0.01, *\*\*\*\*P<* 0.0001. **(B)** CRISPR-Cas9/mClover-LMNA1 mediated HDR assay schematic. **(C)** HDR analyses in WT, RAD51 mutant, and BRCA2 mutant cells. BRCA2 and RAD51 WT and mutant HeLa Kyoto cells were transfected with Lamin A-targeting sgRNA and mClover Lamin A donor constructs before analysis for mClover Lamin A-positive cells (HDR positive cells) after 72 hours. Representative images are shown: DAPI (nuclear DNA), GFP (productive mClover-Lamin A fusion), and merged staining. Scale bar = 50 µm. **(D)** HDR efficiency in WT, RAD51 mutant, and BRCA2 mutant cells. Bar graph showing the mean of HDR positive cells (%) ± SEM from three independent repeats (n>200 per repeat). *P*-values were derived from the one-way ANOVA test, followed by Bonferroni’s multiple comparison test. ns not significant, **** *P*<0.0001.

### TR2i-RAD51 interface mutations regulate RAD51 at stalled DNA replication forks

To further probe these point mutations, we analyzed their role in RAD51 cluster formation and co-localization with single-stranded DNA binding protein RPA upon DNA break damage via high resolution dSTORM (direct Stochastic Optical Reconstruction Microscopy) single-molecule localization microscopy^59^. For this test, we analyzed macromolecular protein clusters for RAD51 and RPA content following neocarzinostatin (NCS) treatment, which induces DNA damage including DSBs (Figures 6A-6C). BRCA2^Tr/Tr^ cells are deficient in assembling RAD51 into larger clusters three hours after NCS treatment when compared to wild-type BRCA2 cells in representative Delaunay triangulations (DT) graphs of RPA and RAD51 molecules. In contrast, cells carrying BRCA2 P3292L, RAD51 M210L and RAD51 M210I mutations initiate damage-induced RAD51 macromolecular assemblies as seen in representative DT graphs (Figures 6A-6C), consistent with HDR proficiency. Some mutant cell clusters appear somewhat smaller compared to those with WT RAD51, suggesting changed RAD51 damage association at some assembly sites. DNA damage including DSB’s also stalls DNA replication forks. As we observed no significant impact of TR2i-RAD51 interface mutations on HDR, we reasoned that cluster differences observed by STORM with genome-wide DNA damage could include RAD51 reactions at DNA replication sites.

**Figure 6.**
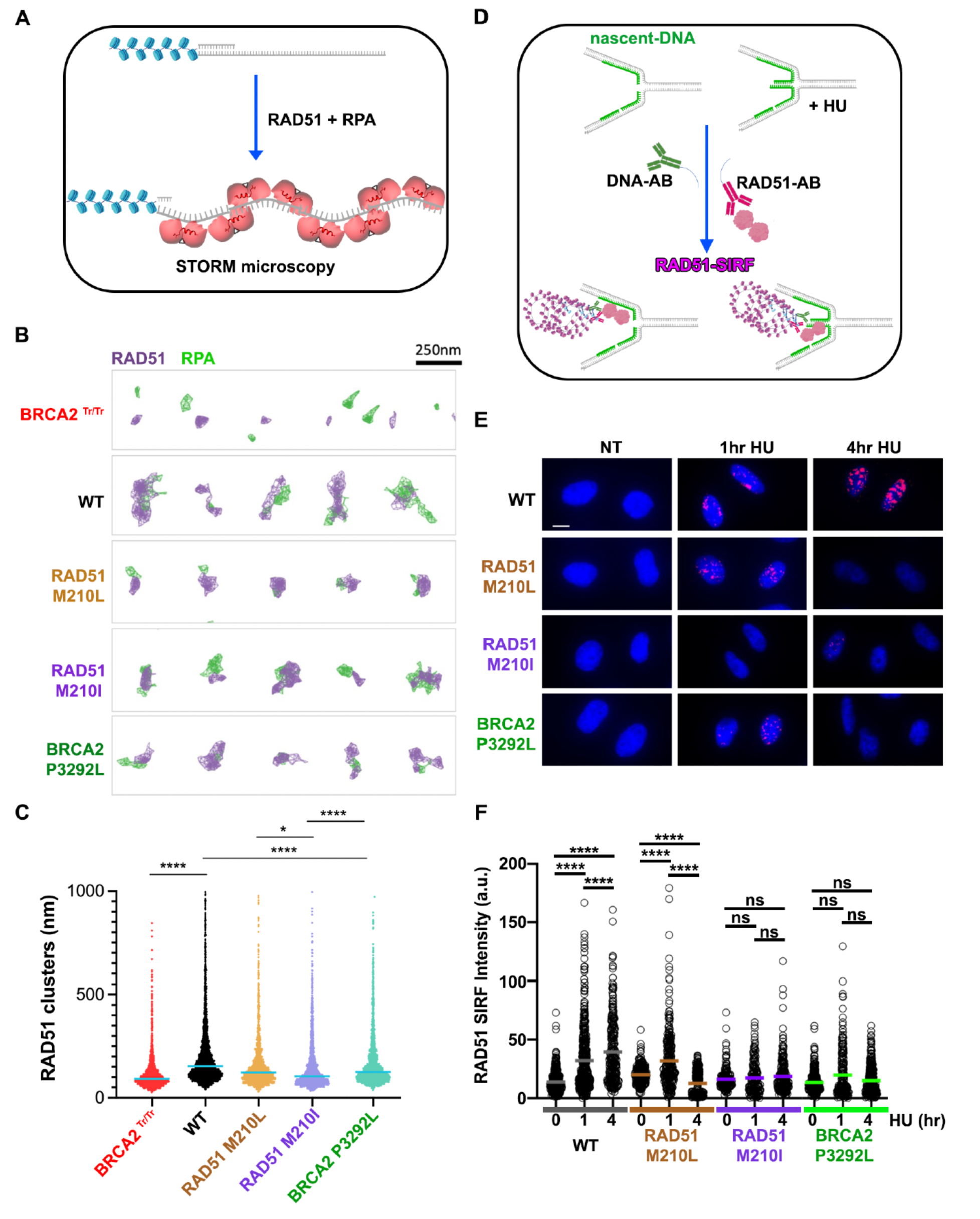
BRCA2 TR2i-RAD51 dimer interaction mutants reveal very different mutant impacts for DNA substrates at replication forks compared to DNA breaks. **(A)** Schematic for recombinogenic RAD51 filaments on ssDNA at breaks. **(B)** Representative dSTORM images (segmented with Delaunay triangulation (DT)) of RAD51 (violet) and RPA (green) cluster in wildtype and mutant HeLa Kyoto cells 3h after exposure to 0.5 µg/ml at Neocarzinostatin (NCS). Scale bar = 250 nm. **(C)** Distribution of observed RAD51 cluster length. *P*-values were derived from the two-sided Wilcoxon rank sum test. \**P*< 0.05, \*\*\**P*< 0.001, \*\*\*\**P*< 0.0001. **(D)** Schematic of the SIRF assay measuring protein association to DNA at nascent replication forks (green DNA) with and without stalling with hydroxyurea (HU, 4 mM). A RAD51-SIRF signal is produced by proximity ligation assay only when antibodies against RAD51 (red) and nascent DNA (green) are within <40nm, indicating protein-DNA fork binding. **(E)** Representative images of RAD51-SIRF with wild-type (WT) and mutant HeLa Kyoto cells disrupting the BRCA2 TR2i interaction with RAD51. Scale bar = 10 µm. **(F)** Scatter Plot of nuclear RAD51-SIRF intensities (a.u. arbitrary units), which measures the association of RAD51 to DNA at nascent replication forks after hydroxyurea (HU, 4 mM) treatment. Bar represents mean, *P*-values were derived from the two-tailed T-test (n = 192-474, three independent biological experiments). ns, not significant; ****, *P*< 0.0001.

To directly assess RAD51 reactions at stalled forks, we determined RAD51-SIRF kinetics, as previously shown to test for RAD51 filament assembly and stability at nascent DNA forks ^60^. Using SIRF (*in situ* protein interaction with nascent DNA replication forks), which measures protein association to newly replicated DNA on a single molecule level ^61^, we find that WT RAD51 accumulates at stalled forks one hour after treatment with hydroxyurea (HU), and continues to associate with stalled forks four hours after HU treatment, indicating stable RAD51 associations at stalled forks in wild-type cells (Figures 6D-6F and Figures S5B-S5D). In contrast, cells containing the least disruptive RAD51 M210L mutant show a modest initial increase in RAD51 at stalled forks, and a sharp reduction four hours after HU treatment (Figures 6D-6F, beige, and Figures S5B-S5D). These results suggest that initial RAD51 loading is less efficient and RAD51 stability at stalled forks is compromised in these mutant cells. Most prominently, RAD51 M210I fails to significantly associate with stalled fork at early or late timepoints following HU treatment (Figures 6D-6F, beige, and Figures S5B-S5D). These data suggest that in contrast to DSB sites where RAD51 M210I robustly assembles into break-associated damage foci sufficient for HDR (Figures 5C and 6B), the mutant RAD51 fails to bind stalled forks. This highlights the functional importance of RAD51 M210 for FP. BRCA2 P3292L mutant cells follow this same trend albeit initial binding defects are more noticeable (Figures 6D-6F, green). Collective findings suggest that the requirements for RAD51 loading at stalled DNA forks differ from those at DNA breaks, whereby TR2-RAD51 is required for the former.

## DISCUSSION

BRCA2 controls RAD51 in both HDR and FP to suppress genome instability associated with cancer. So the conceptual understanding has been that these two functions were interrelated with BRCA2 promoting: 1) RAD51 loading onto RPA-coated ssDNA essential for HDR, and 2) subsequent RAD51 filament stabilization, as less important to HDR but essential for FP. Yet, BRCA2-mediated HDR and FP are physiologically and genetically separable ^6,8^. Here collective findings reveal BRCA2 accomplishes its separate function at stalled replication forks by clamping RAD51 into a unique dimer conformation opening the DNA binding channel and so favoring RAD51 dimer binding to B-form dsDNA and short ssDNA that predominate at stalled and reversed forks ^62,63^. Long ssDNA regions and non-B-DNA structures on the other hand are limited at forks to prevent DNA replication instability. Notably, recent thought-provoking biochemical data suggest that prevention of MRE11-dependent DNA degradation requires protection by TR2i-RAD51 stabilization on dsDNA rather than ssDNA ^25^, in accord with the findings presented here. Importantly, the crystal structure of TR2i with RAD51-dimer is in the absence of DNA, i.e. prior to binding DNA. Thus, our results delineate how BRCA2 TR2i reshapes RAD51 dimer conformation to promote RAD51-dsDNA binding that supports BRCA2 C-terminal FP roles beyond RAD51 filament stabilization: BRCA2 TR2 dictates DNA substrate specificity for RAD51 nucleation onto dsDNA.

A RAD51 dimer is the fundamental building block for RAD51 assembly onto DNA^21,64^. During HDR, RAD51-ATP dimers assemble onto ssDNA in a conformation that extends the ssDNA. This forms a stable nucleoprotein complex that surface-exposes DNA bases, and so enables the search for base pair homology ^17,24,29,65^ (Figure 7, left). The BRCA2 TR2i clamp allosterically reshapes the RAD51 dimers into dimers suitable for RAD51 nucleation on B-DNA and dsDNA in the presence of ATP. So far it was thought that BRCA2 loads RAD51 onto ssDNA and subsequently stabilizes a filament already assembled onto DNA. Our collective data supports an additional model to this canonical inter-related two-step model: TR2i-RAD51 complexes are formed prior to nucleation enabling RAD51 binding to dsDNA, a reaction that is principally inhibited by BRC repeats. This concept predicts that FP and HDR are distinct reactions defined by opposing and perhaps mutually exclusive DNA substrates, whereby BRCA2 TR2i promotes RAD51-ATP loading and stability on non-extended, B-form and dsDNA fork substrates (Fig 7, right).

For HDR, BRCA2 repeats BRC1-4 assemble RAD51 protomers while BRC5-8 stabilize RAD51 protomers on ssDNA, together enabling RAD51 assembly and filament growth on longer ssDNA as suited for HDR ^24,25,29^. Yet, BRC4 promotes RAD51 disassembly from dsDNA ^24,25,29–32^. All these activities derive from two distinct consensus sequence interface motifs (residues FxxA and LFDE) in the BRC repeat interaction with RAD51 ^34^. Importantly, the FxxA motif in the BRC repeat competes with this same sequence motif that is present in the PM of RAD51, which promotes protomer oligomerization. In contrast, we find that the RAD51 FxxA PM linchpin for the RAD51 dimer ^35,44,47,66^ is functionally stabilized by BRCA2 TR2i (S3291:M210:F86 triad). Indeed, our results show that the BRCA2 BRC repeats not only compete for the buried phenylalanine binding site, but also clash with the critical TR2i tight turn (Figures S6A and S6B), whereas TR2i stabilizes the RAD51 dimer. These combined observations provide a compelling structural basis for an antagonistic rather than sequential relationship between RAD51 loading on ssDNA for HDR and RAD51-B-DNA binding and stability for FP.

Our findings establish RAD51 M210 as a key RAD51-identifying residue crucial for BRCA2 TR2i recognition. Furthermore, they unveil how sites previously implicated in BRCA2 function (BRCA2 S3291, P3292, F3298) form structural components that allosterically control RAD51 dimer conformation. BRCA2 TR2 F3298 acts as a latch over RAD51 G155, which is part of the main chain linking to the ATP binding site. The structural importance of this site is consistent with an allosteric change of this site required for reshaping into an asymmetric RAD51 dimer. RAD51 M210 hydrogen bonds to BRCA2 S3291. Cell cycle specific S3291 phosphorylation efficiently regulates this interaction as the bulky negatively charged phosphate group geometrically clashes with M210 side chain in a strong sterically-driven switch. Serine to Alanine mutation also removes this sulfur hydrogen bond in the S3291:M210:F86 residue triad, thus weakening the complex despite not changing the charge. Interestingly, instead of Methionine, the meiosis-specific RAD51 homologue DMC1 holds a Lysine at the analogous site (K209). The flexibility of the Lysine and its positive charge could accommodate S3291 phosphorylation, explaining the preserved interaction of the phosphorylation motif with DMC1 ^67^ (Figure S6C). Of note, the S3291 is buried within the TR2-RAD51 dimer, suggesting that CDK mediated phosphorylation must occur before TR2 binding to RAD51. This makes it unlikely that CDK controls RAD51 stability after RAD51 filament assembly as has been postulated so far, thus further strengthening the importance of the TR2-RAD51 interaction and its control prior to DNA binding.

The structural significance for M210 in the TR2i interface to fork reactions is underscored by cellular functional assays. Cells with either BRCA2 P3292, RAD51 M210L or M210I mutations remain proficient for RAD51 DNA damage foci assembly and HDR. In contrast, these cells are defective in FP, with unstable and inefficiently bound RAD51 at stalled forks. Most strikingly, RAD51 M210I shows an inability to load onto stalled forks, coinciding with the FP defect; yet, it can load onto DNA damage sites and efficiently repair DSBs. Thus, our findings on M210I reveals a pivotal difference between RAD51 DNA substrates at forks and breaks, which is highlighted when comparing fork and break specific cellular assays. Notably, M210I is a cancer somatic mutation with so far missing clinical annotation ^54^, whereby our analyses here show a loss of genome stability function supporting cancer association.

Recent cryo-electron microscopy (cryo-EM) data show TR2-RAD51-filament complexes with DNA ^45^. While these data support the TR2i architectural arrangement at the RAD51 dimer interface shown in our crystal structure, the cryo-EM analysis proposed an electrostatic interface centered on a RAD51 acidic patch in the DNA-BRCA2 TR2-RAD51-filament complex. Based on our structure and electron density maps that revealed the detailed atomic structure of this area, the solvent-exposed RAD51 acidic patch would be expected to predominantly affect BRC-RAD51 interactions and is suboptimal for interactions with BRCA2 TR2, consistent with the experimental data shown in the cryo-EM report. Extreme C-terminal mutations affect RAD51 interaction and HDR in Ustilago maydis^68^, which is a fungus that contains a single BRC repeat and lacks the additional BRC repeats seen in higher eukaryotes that evolved to load and stabilize RAD51-ssDNA filaments. In analogy, when artificially deleting human BRC 5-8 repeats, which otherwise stabilize RAD51 on ssDNA, TR2 mutations exhibit a mild HDR defect^39^. These results explain how the weaker acidic patch interactions of TR2 with RAD51 can partially, albeit suboptimally, compensate for the missing BRC repeats during HDR. However, mutations in TR2 do not affect HDR proficiency in any human cell system or mouse system tested provided that BRC 1-8 are present, suggesting that TR2 plays no role during HDR under biologically relevant conditions. In contrast, TR2 mutations abrogate B-DNA binding and FP. The duplication of BRC repeats during evolution and concomitant irrelevance of TR2 for HDR underscores the possibility that HDR and FP evolved to become spatially and functionally separate pathways.

Importantly, our detailed structure of the TR2i-RAD51 dimer complex crystalized without DNA. This enabled identification of the pivotal role for M210 in recognizing S3291 within TR2i triad anchors that allosterically reshape RAD51 dimer prior to DNA binding. This was not considered so far due to the canonical view of RAD51 function on ssDNA during HDR, which is well established to be accomplished by BRC repeats. Furthermore, BRC repeats dissemble RAD51 from dsDNA further cementing the notion that RAD51’s biological function is accomplished via ssDNA interactions, leading to the corollary accepted belief that FP is a form of HDR at replication forks. Instead, the crystal structure in the absence of DNA reveals that TR2 reshapes RAD51 prior to DNA binding to promote rather than inhibit B-form DNA binding. These insights change our understanding of BRCA2 C-terminal function from RAD51-ssDNA filament stabilization to RAD51 loading on distinct B-form DNA substrates, suggesting TR2 may control pathway choice between FP and HDR as mutually exclusive reactions. Collectively, our findings identify TR2 as an allosteric control for distinct RAD51 dimer conformations that mediate BRCA2 replication fork stability functions.

## SUPPLEMENTAL INFORMATION

Figures S1 to S6

Tables S1 to S3

Methods

## ACKNOWLEDGMENTS

We thank Sunetra Roy (Department of Cancer Biology, UTMDACC) for critical feedback to improve the manuscript. We acknowledge Dr. Kalina Haas for her inputs on dSTORM cluster data analysis. We acknowledge Marko Hyvonen, Luca Pellegrini (Department of Biochemistry, University of Cambridge) for early studies on dimeric Pyrococcus RadA and TR2, and Sir Tom Blundell (Department of Biochemistry, University of Cambridge). This research was supported in part by the National Institute of Health (NIH) R35 CA220430 to J.A.T., P01 CA092584 to J.A.T., 1R01ES029680 to K.S., F31 CA142313 to M.A.L, Cancer Prevention and Research Institute of Texas (RP180813 to K.S. and J.A.T., RP180463 to K.S.) and Robert A. Welch Chemistry Chair (J.A.T.), K.S. and J.A.T. are CPRIT scholars in Cancer Biology. K.S. is a Rita Allen Foundation Fellow. Work in A.R.V.’s laboratory was supported by UK Medical Research Council Program Grants MC_UU_12022/1 and MC_UU_12022/8, Gray Foundation Team Science Awards 235599 and 281760.5119253.0409, and an MOE Academic Research Tier 3 (RIE2025) MOE-MOET32021-0004 from the Ministry of Education, Singapore. Beamlines 8.3.1 and 12.3.1 at the Advanced Light Source are operated under support from NIH (P30 GM124169) and the Integrated Diffraction Analysis Technologies Program of the US Department of Energy Office of Biological and Environmental Research.

## AUTHOR CONTRIBUTIONS

M.A.L., K.S. and J.A.T. conceptualized the project. K.S. and J.A.T directed the study. M.A.L., Z.W.Z. J.A.T., A.R.V. and K.S. contributed to design, interpretation, and supervision for experiments. M.A.L, A.A, R.L.P., S.N, X.W., and C.L.T did structural, biochemical, and in vitro binding experiments. S.M.A, M.L and L.R.K generated the CRISPR mutant Hela Kyoto cells. S.M.A. performed DNA fiber experiments, cellular survival assay, HDR and S.M.A., W.E. and W.S. performed STORM experiments, Y.C. performed SIRF studies. M.A.L, J.A.T. and K.S. wrote the manuscript, and K.S., M.A.L, J.A.T., A.R.V and C.L.T. reviewed and edited the manuscript.

## INCLUSION AND DIVERSITY

We support inclusive, diverse, and equitable conduct of research.

Data and materials availability: The structure has been deposited in the Protein Data Bank under accession code 8UVW. Hela Kyoto cells are available upon request to A.R.V. All data are available in the main text or the supplementary materials.

---

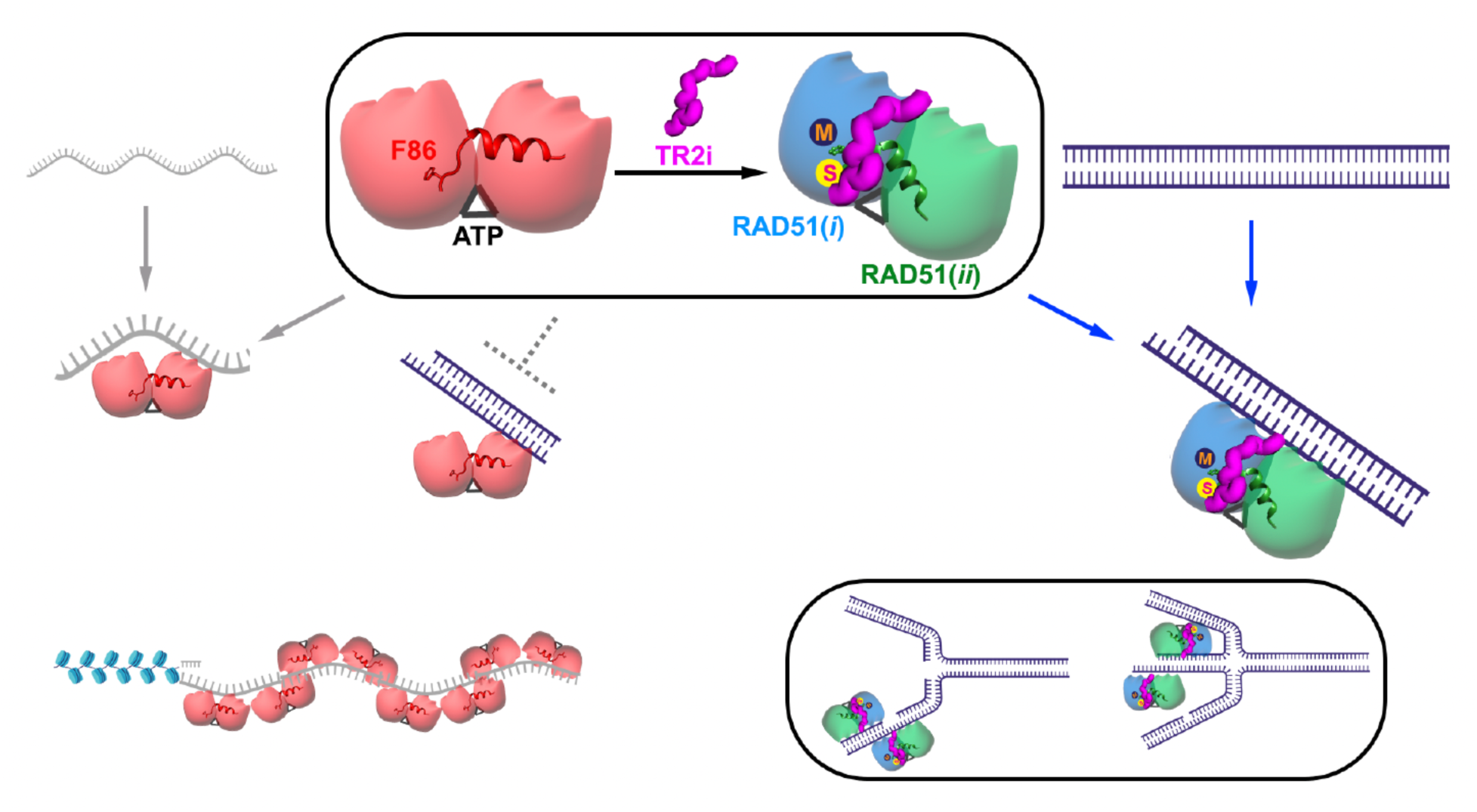
A model for BRCA2 FP in which TR2i binding reshapes ATP-bound RAD51 dimer to efficiently bind short dsDNA as suitable for FP but not HDR. A RAD51 dimer in the ATP-extended conformation (red) readily binds flexible ssDNA (grey arrows, far left) at processed DNA breaks (bottom left) but is inefficient in binding duplex B-DNA (grey dashed line). Upon BRCA2 TR2i binding (magenta), the TR2i-clamped RAD51 dimer (blue and green subunits) with S3291 and M210 interactions (M,S spheres) is reshaped for optimal binding to B-form DNA (blue arrows, far right), which predominates at stalled and reversed DNA replication fork substrates (bottom right).

