## supplemental figures tables and methods for "BRCA2 C-terminal clamp restructures RAD51 dimers to bind B-DNA for replication fork stability"

### SUPPLEMENTAL INFORMATION

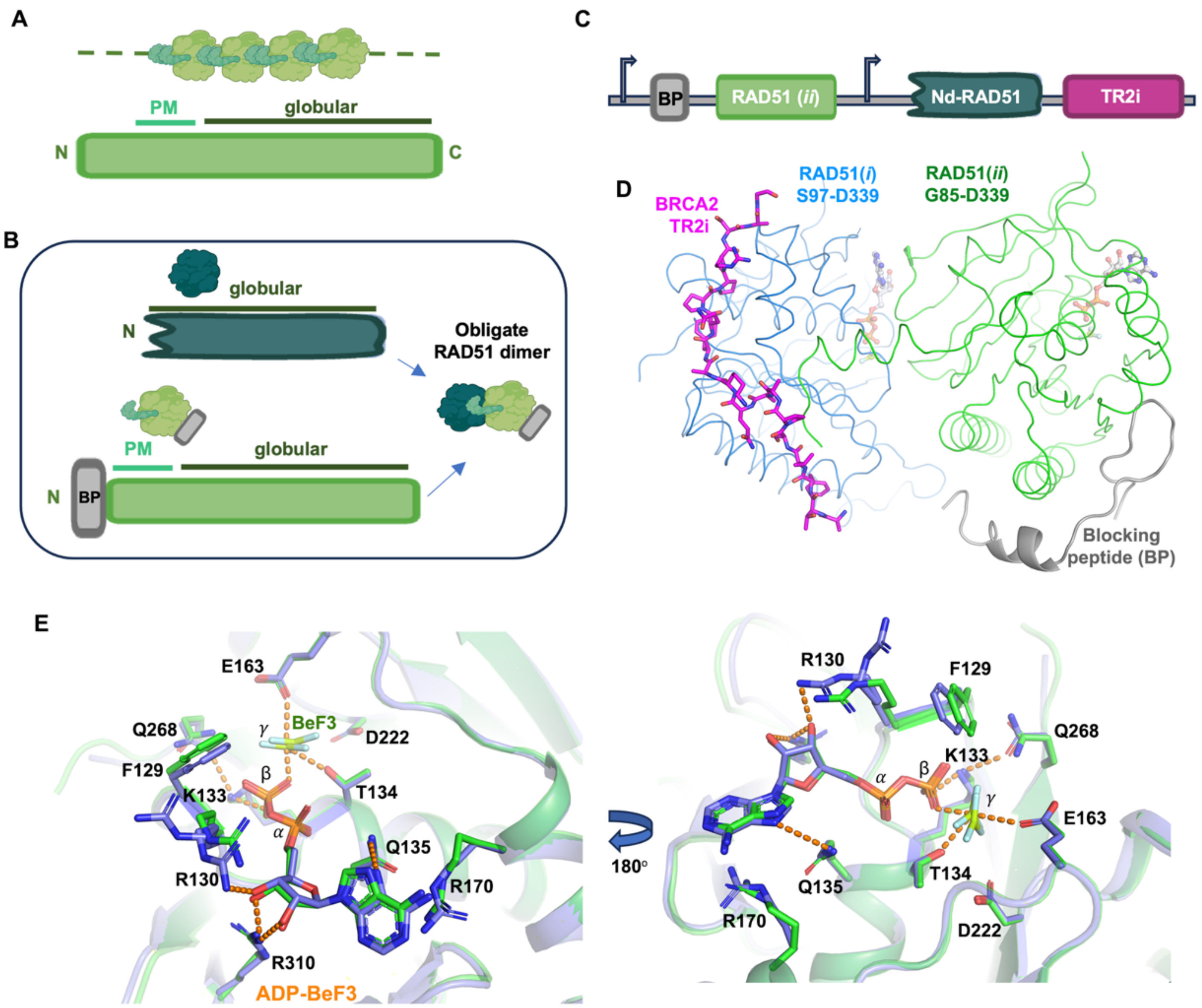


**Figure S1. Crystal structure of BRCA2 TR2i-RAD51 dimer complex, related to Figure 1**

**(A)** RAD51 gene schematic containing N-terminal FxxA polymerization motif (PM) and globular (C-terminal) ATPase interaction domain.

**(B)** Sketch of RAD51 (*i*) and RAD51 (*ii*) construct design forming an obligate dimer without possibility of oligomerization in either N-terminal or C-terminal direction. N-terminal domain and oligomerization motif (FxxA) of RAD51(*i*) are removed (S97-D339, blue). RAD51(*ii*) (G85-D339, green) retains the FxxA oligomerization motif, enabling RAD51*(i)* - RAD51*(ii)* dimer formation, but contains a polymerization blocking peptide preventing polymerization in the C-terminal/reverse direction and thus larger RAD51 complexes.

**(C)** Crystallization expression construct of TR2i-RAD51 dimer complex. BP, polymerization Blocking Peptide; L, flexible linker, RAD51*(ii),* RAD51 G85-D339 (green); RAD51*(i),* RAD51 S97-D339 (blue); TR2i, BRCA2 amino acids 3289-3303 (magenta).

**(D)** BRCA2 TR2i structure (magenta sticks) at the RAD51 dimer interface (subunit fold as blue and green ribbons) with blocking peptide (BP, grey). ATP-transition state analog ADP-$BeF_3$ is present in both RAD51 protomers (pale sticks, center and right).

**(E)** Overlay of two ADP-BeF3 binding sites (from each protomer). ATP binding sites overlap showing that TR2i binding does not substantially alter the ATP binding site geometry between the RAD51 protomer subunits.

F129, R130, and R170 display some confirmational variability in the shared ATPase interface.

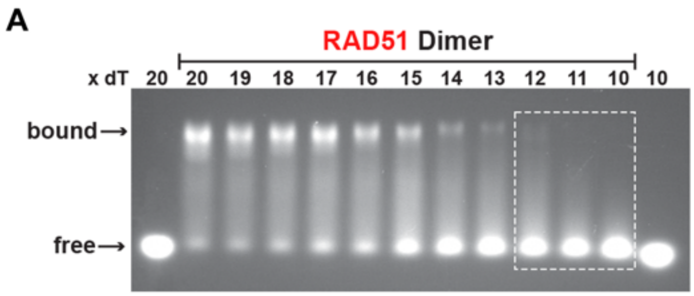


**Figure S2. BRCA2 TR2i-RAD51 dimer binds short B-form DNA including short ssDNA, related to Figure 2**
**(A**) Electrophoretic mobility DNA shift assays (EMSA) of RAD51 binding short ssDNA varying in DNA lengths (10-20 dT).
**(B**) EMSA of TR2-RAD51 binding short ssDNA varying in DNA lengths (10-20 dT) suggests that TR2 allows RAD51 binding of hyper-short ssDNA consistent with base stacked or B-DNA rather than an extended DNA conformation.

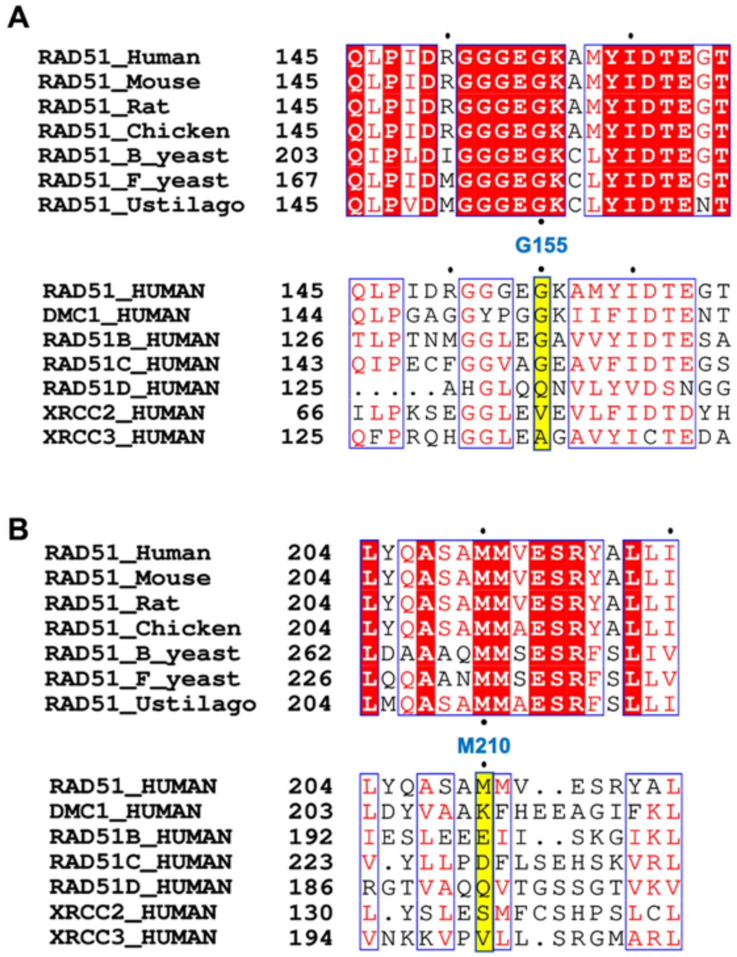

**Figure S3. Proline driven TR2i-RAD51 interface triad residues are conserved in RAD51 but not paralogs, related to Figure 3**
**(A)** Sequence alignment of RAD51 homologs with conserved G155 (top), and moderate conservation with RAD51 paralogs (bottom). **(B)** Sequence alignment of RAD51 homologs shows conservation of M210 amongst species (top), contrasting with the poor conservation of the methionine at this position when comparing the amino acids with RAD51 paralogs (bottom), and supporting specific importance of M210 for RAD51.

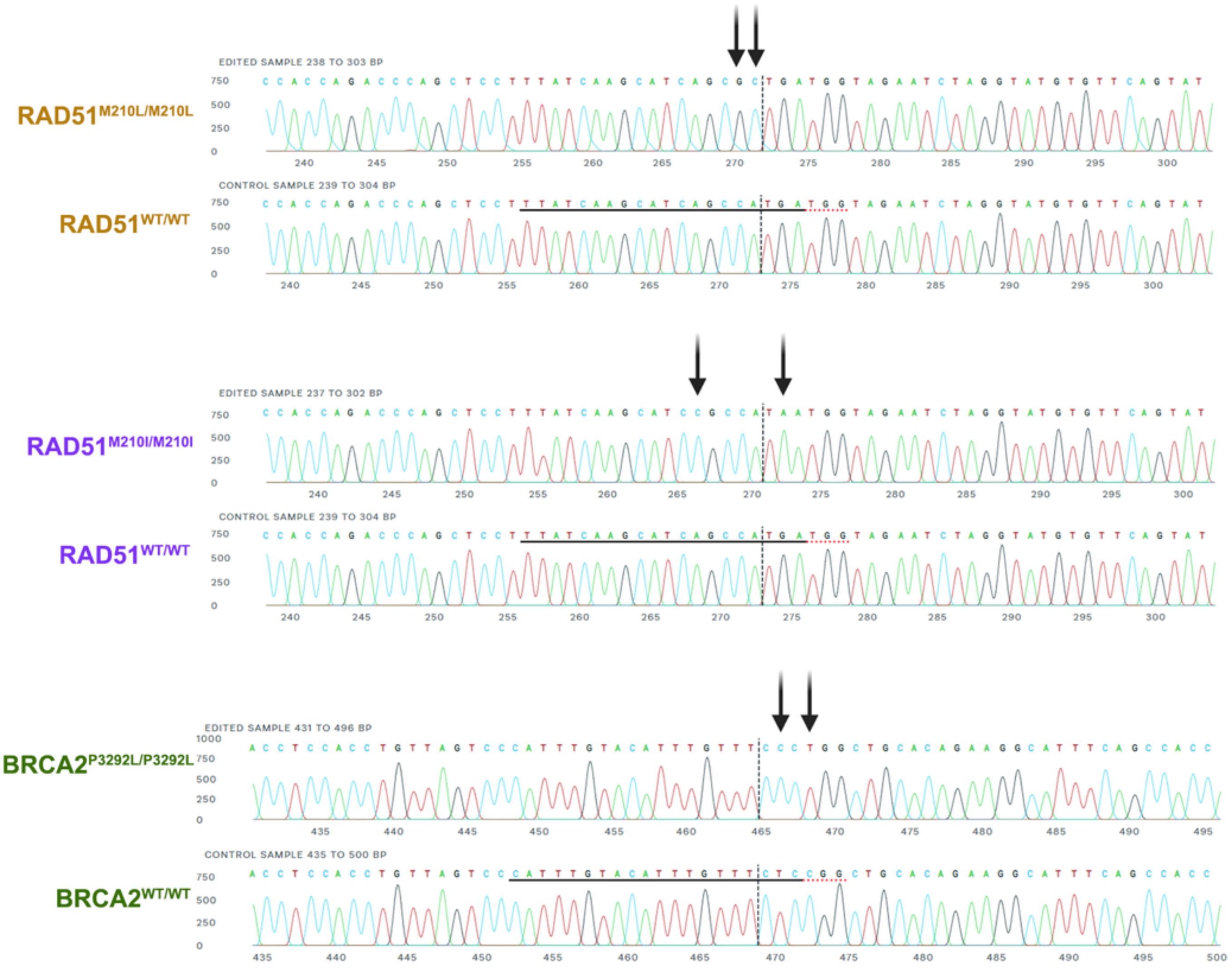


**Figure S4. CRISPR-Cas9 mutation confirmation of M210L/M210I RAD51 and of P3292L BRCA2 mutations by Sanger sequencing from single cell colony, related to Figures 5-6**

Existence of the targeted knock-in mutation was checked for each isolated clone by Sanger sequencing after PCR amplification of the target region. Successful homozygous knock-in mutants were obtained and used in this study.

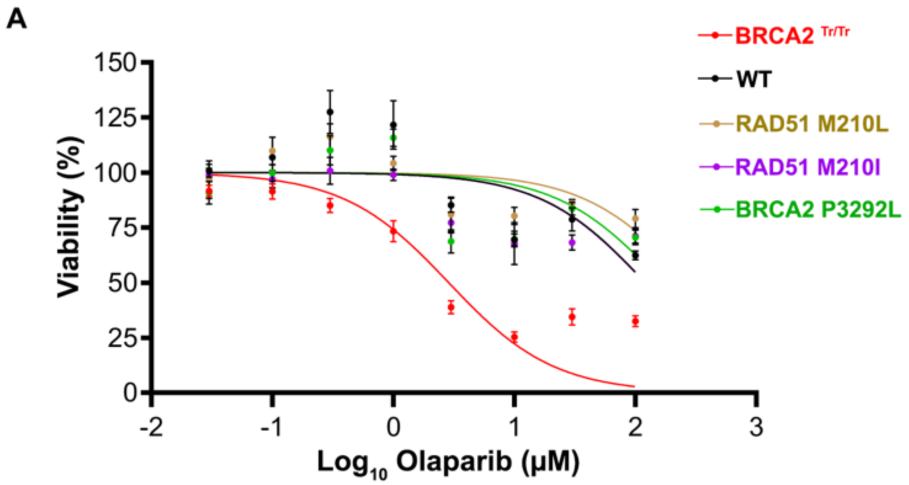


| Cell Line | Log $IC_{50}$ (mean ± SD) |
|---|---|
| BRCA2 Tr/Tr | 0.4822 ± 0.194 |
| WT | 2.080 ± 0.060 |
| BRCA2 M210L | 2.503 ± 0.414 |
| BRCA2 M210I | 1.927 ± 0.695 |
| BRCA2 P3292L | 2.231 ± 0.263 |

*** ns ns ns

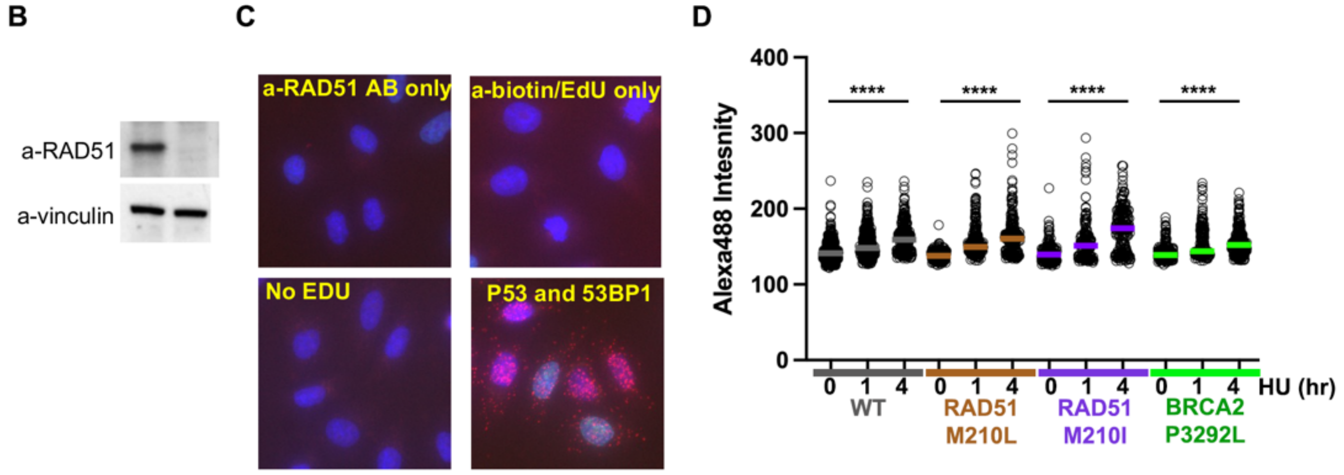


**Figure S5. PARP inhibitor sensitivity and RAD51-SIRF assays in TR2i-RAD51 mutant cells, related to Figures 5-6**

**(A**) Olaparib dose response curves (*top*) for WT and mutant HeLa Kyoto cells after exposure to the indicated dose of olaparib for 72 hrs. Error bars represent SEM from triplicate experiments. Mean IC50 ± SD for each genotype is shown (*bottom*). Unpaired student's t-test was used for statistical analysis. ns, not significant; ∗∗∗, *P*< 0.001.

**(B)** RAD51 knock down by siRNA in U2OS cells confirms detection of RAD51 by the antibody used in the RAD51 SIRF assays.

**(C)** Omission of either primary anti-RAD51 antibody or the anti-biotin antibody (which detects the nascently EdU labeled DNA) does not produce a RAD51 SIRF signal, nor do reactions where EdU is omitted in the cell culture, confirming specificity of the assay to detect RAD51 protein binding to nascently replicated DNA. A p53-53BP1 PLA serves as positive control for the PLA reaction.

**(D)** Click-IT with Alexa488 azide show that the amounts of EdU incorporated into nascent DNA overall show the same trend between the mutant (BRCA2 P3292L, green; RAD51 M210L, brown; RAD51 M210I, purple) and WT (gray color) cell lines with Alexa488 intensities slightly increasing over time likely due to residual EdU incorporation, ruling out that the RAD51 SIRF changes are caused by limiting EdU available for the assay reaction. ∗∗∗∗, *P*< 0.0001

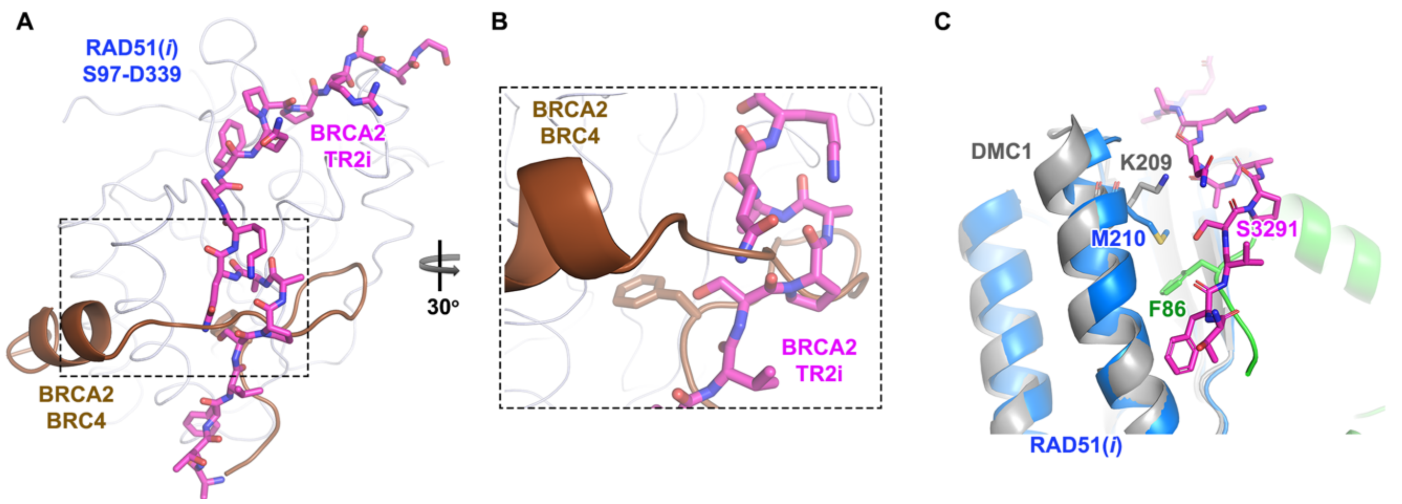


**Figure S6. BRCA2 TR2i geometrically opposes BRCA2 BRC4-RAD51 interaction, related to Discussion**
**(A)** BRCA2-TR2i (magenta carbons) and BRC4 (brown ribbon) overlay from superimposed RAD51 protomers (pale tubes) reveals a perpendicular spatial relationship, geometrically clashing by crossing in proximity to S3291. BRCA2 BRC4 motif (brown tube) clashes in space with the critical S3291:M210:F86 residue triad interface (magenta carbons). Such clash explains the structural basis for the antagonistic relationship of the two distinct BRCA2:RAD51 interactions.
**(B)** Close-up view. Hairpin of BRC4 (brown tube) clashes with the TR2i tight-turn motif (magenta carbons
**(C)** Superposition of DMC1 (gray) and RAD51(*i*) (blue) structures displaying the RAD51 M210 position (blue carbons) is equivalent to DMC1 K209 (gray carbons, PDB: 1V5W). M210 sulfur stacks with RAD51(*ii*) (green) F86 from the polymerization motif.

**Table S1. X-ray data collection and refinement statistics**

| | **RAD51-BRCA2 Cter** |
|---|---|
| Wavelength (Å) | 0.9795 |
| Space group | $P2_12_12_1$ |
| Cell dimensions | |
| *a*, *b*, *c* (Å) | 51.46, 98.55, 128.32 |
| α, β, γ (°) | 90, 90, 90 |
| Resolution (Å)[a] | 39.24 – 2.73 (2.86 – 2.73) |
| Observations[a] | 228267 (31885) |
| Unique observation[a] | 17974 (2352) |
| $R_{pim}$[a] | 0.041 (1.794) |
| Mean $I/\sigma I$ [a] | 15.9 (0.6) |
| Completeness (%)[a] | 99.8 (99.9) |
| Multiplicity[a] | 12.7 (13.6) |
| CC(1/2)[a] | 0.999 (0.340) |
| Refinement | |
| $R_{work}/R_{free}$ (%) | 24.08/28.55 |
| No. of atoms | |
| Protein | 4033 |
| Ligands | 70 |
| Water | 30 |
| rmsd bond length (Å) | 0.003 |
| rmsd Bond angles (°) | 0.651 |
| Average B-factor | 104.09 |
| Protein | 104.29 |
| Solvent | 94.69 |
| Ligands | 96.75 |
| Ramachandran (%) | |
| Favored | 94.38 |
| Allowed | 5.22 |
| Outliers | 0.40 |
| PDB code | **8UVW** |

[a] Values in parentheses are the statistics for the highest resolution shell of data.

| Table S2. Oligo used in HDR assay | | |
|---|---|---|
| **Desired Mutation** | **HDR Template** | **Diagnostic Restriction Site** |
| M210L | CTCTCTGGCAGTGATGTCCTGGATAATGTAGCATATG CTCGAGCGTTCAACACAGACCAC CAGACCCAGCTCCTTTATCAAGCATCAGCGCTGATGG TAGAATCTAGGTATGTGTTCAGT ATAAGACACCAAATATGTTCTTAAGAGTCCTTCCCTGA ATCTTGTAATGGCTATTTGGCC | HaeII |
| M210I | CTCTCTGGCAGTGATGTCCTGGATAATGTAGCATATG CTCGAGCGTTCAACACAGACCAC CAGACCCAGCTCCTTTATCAAGCATCCGCCATAATGG TAGAATCTAGGTATGTGTTCAGT ATAAGACACCAAATATGTTCTTAAGAGTCCTTCCCTGA ATCTTGTAATGGCTATTTGGCC | FokI |
| P3292L | CATCTGAGGAGAATTCAGTTCTTTTTTCTTTATGGGTG TTTCGTATTTGGTGCCACAACTCCTTGGTGGCTGAAAT GCCTTCTGTGCAGCCAGGGAAACAAATGTACAAATGG GACTAACAGGTGGAGGTAAAGGCAGTCTACTCAAGAA ATCCAAGGCTCTTCTCTTTTTGCAGTTCTT | BstNI |

| Table S3: Cell lines used in this study | | |
|---|---|---|
| **Cell Line** | **Description** | **Targeted Domain** |
| HeLa Kyoto | HeLa Kyoto cell line | - |
| HeLa BRCA2 KO | BRCA2 KO using CRISPR-Cas9 System | BRCA2 Exon 11 |
| HeLa RAD51 M210L | HeLa with M210L homozygous knock in | RAD51 Exon 7 |
| HeLa RAD51 M210I | HeLa with M210L homozygous knock in | RAD51 Exon 7 |
| HeLa BRCA2 P3292L | HeLa with P3292L homozygous knock in | BRCA2 Exon 27 |

### Materials and Methods

#### Crystallization constructs: cloning, expression, and purification

For structural characterization, two RAD51 subunits were cloned into a pRSFduet1 (Novagen) plasmid modified with a N-terminal 6Xhis tagged Maltose Binding Protein domain in ORF1 (Nco1-Asc1). RAD51gene encoding sequence G85-D339 (RAD51-*ii*) fused with a (3X Thr-Gly-Ser) linker to a RAD51-polymerization blocking peptide (BRC4 K1517-D1547) at its N-terminus, was cloned into ORF1 (between Asc1 and Not1) in frame with 6XhisMBP, with an intervening TEV protease cleavage site (fig. S1A). The pRSFduet ORF2 contains RAD51 gene encoding sequence S97-D339 (RAD51-*i*) fused with a minimized Oct4 linker (SSDSSLSSPS) and a BRCA2 TR2i peptide (A3270-G3305) at its C-terminus, was cloned between NdeI and AvrII (ORF2). Each ORF CDS encoding a RAD51 subunit contain alternative codons to prevent recombination between ORFs. The RAD51(*i*) contains S208E and A209D mutations, which prevent BRC4 binding during co-expression. Both RAD51(*i*) and (*ii*) contain C319S mutation, which improved crystal quality. The resulting construct (TR2i-RAD51 Dimer) expresses an engineer, obligate dimeric RAD51 with a stably bound BRCA2 TR2i interaction. The RAD51(*i)* protomer (RAD51 S97-D339) without BRCA2 TR2i fusion produced dimer (RAD51 Dimer) alone.

The TR2i-RAD51 Dimer construct was transformed into Krome (DE3) expression strain (2X StrepII-tag incorporated into genomic dnak gene) (Poseidon Laboratory) containing pRARE2 plasmid and plated on LB supplemented with antibiotics kanamycin and chloramphenicol. The starter culture was scraped directly from plated colonies and grown at 37°C for ~ 5 hrs in Turbo broth expression media supplemented with kanamycin & chloramphenicol before being inoculated in large scale culture at 37°C for ~ 5 hrs until OD ~ 2.0. Cultures were cooled to 30°C before induction with 200 μM IPTG, followed by overnight expression at 15°C for 16 hrs. Cells were harvested by centrifugation at 5000 *g*, 4°C for 30 min and resuspended in 20 ml of 20 mM Tris pH 8.0, 500 mM NaCl, 20 mM Imidazole supplemented with EDTA-free protease inhibitor tablets (Sigma) and flash frozen with liquid N2.

The cells were lyzed in an emulsiflex C-5 (avestin) high-pressure homogenizer with the addition of 1 mM TCEP. The TR2i-RAD51 Dimer complex was purified by AKTA on a 25 ml of Ni-NTA column with buffer A (Tris pH 8.0 and 500 mM NaCl) and eluted with buffer B (Tris pH 8.0, 500 mM NaCl, and 200 mM Imidazole, pH 8.0). Eluate was further purified over 15 ml of amylose resin (NEB) by gravity, washed with buffer A, and eluted by in-column TEV protease digestion. Basically, 1:100 (w/w) tev protease was added to the washed amylose resin in column, which was capped and resuspended for 2 hrs at RT with rotation to maintain suspension. The resin was settled at 4°C after digestion, and the protein was eluted by gravity with buffer A. Fractions containing sample were concentrated to ~ 4 ml, supplemented with imidazole to 10 mM, and passed through 10 ml of Ni-NTA columns to remove his-tagged Tev protease and any uncleaved sample, and 10 ml of streptactin resin to remove minor contaminating DNAK bound-fraction of sample. Protein sample was then polished by Superdex 200 (26/100) column with buffer C (10 mM Tris pH 8.0 and 300 KCl). The purified TR2i-RAD51 Dimer complex was concentrated to 1.45 mM, and flash frozen in liquid N2.

RAD51 Dimer (without TR2i peptide) over-expression culture and purification was performed largely as described above with some exceptions. Dimer construct was transformed into Rosetta 2 expression strain (Novagen). The initial capture step was done with a 50 ml of Ni-NTA XK 50/20 column, and washed and eluted as before. The pooled fractions were spiked to 5 mM TCEP. The subsequent amylose/TEV cleavage was performed with a 20 ml of amylose resin. TEV cleavage and elution procedures were performed as before. The clean-up steps with Ni-NTA were performed as before, but the dimer sample was spiked with 5 mM TCEP.

For size exclusion chromatography, the dimer sample (in 5 mM TCEP) was concentrated to ~4 ml, and purified through a Superdex 75 (26/100) column equilibrated in 20 mM Tris, 500 mM NaCl, and 1 mM TCEP. The fractions corresponding to the dimer peaks were further polished and buffer exchanged using Superdex 75 (16/60) equilibrated in 20 mM Tris, 150 mM KCl, and 1 mM TCEP. Stoichiometric RAD51 Dimer was concentrated, and flash frozen for storage at -80°C

**Protein crystallization**

TR2i-RAD51 Dimer complex (315 μM, ~20 mg/ml) was prepared with 1.2 molar excess 11dT DNA (dissolved in 10 mM Tris pH8.0 and 100 mM KCl) in the presence of 2.5 mM ADP-$BeF_3$. Final complex was brought to 5 mM $MgCl_2$. Sitting drops were set up in 1:2 drop ratio of protein:crystallization buffer condition containing 50 mM MOPS pH 7, 25% Dioxane, 5 mM $MgCl_2$ and 1 mM Spermine at 20°C. Crystals grew to maximum size after 2 weeks, and were cryoprotected in crystallization buffer plus 10% glycerol before being frozen in liquid nitrogen. Diffraction was improved by mutating TR2i Cys319 residue to Ser, which corresponded to the RAD51 homolog DMC1 residue (C319S).

**Data collection, structure determination, and structure analysis**

The diffraction data were collected at 0.9795 Å at beamline 12-1 at the Stanford Synchrotron Radiation Lightsource (SSRL) in Menlo Park, CA. The collected dataset was integrated using the HKL2000 program [69] and scaled and merged in the CCP4 suite using aimless software [70]. The phase was solved via molecular replacement with the human RAD51 structural model (PDB: 1N0W) in Phenix using Phaser [71]. The asymmetric unit contained two RAD51 subunits, and the peptides of BRCA2 TR2i and BRC4 (a blocking peptide) as well as the linkers were manually built using Coot software [72]. The final model was refined using Phenix.refine to $R_{\mathrm{work}}$ = 0.24 and $R_{\mathrm{free}}$ = 0.28. Although 11dT ssDNA was present in the crystallization solution, unstructured DNA-binding loop regions and electron-density maps reveal a DNA-free structure.

Structural alignments and analysis of RAD51:RAD51 interface orientations were performed in PyMol. Protomer *ii* of the TR2i-RAD51 dimer was superimposed against equivalent subunits of RAD51 dimers (alpha-carbons, 200 atoms, RMSD = 0.5) extracted from published multimeric structures (PDB: 5NWL-chain A&B, E&F, 8BSC). The orientation of the corresponding RAD51 protomer *i* was compared to that of the equivalent subunit in various nucleotide bound states and conformations. Movies were made by morphing TR2i-RAD51(*i*) with extended or compressed ATP-RAD51 subunits using Chimera.

**Pulldown construct: cloning, expression, and purification**

The three pull-down constructs were individually cloned into the pRSFduet vector. The resulting three open reading frame (ORF) constructs contain 1) an ORF encoding for expression of full length wt-RAD51 or point mutants (between NcoI and SacI), 2) and an ORF encoding for an N-terminal MBP-TR2i peptide (A3270-K3314)-8XHis tag, and 3) an ORF encoding an N-terminal MBP-tagged-RAD51 ATPase domain (H93-D339 with M210L mutation)-StrepII tag to control RAD51-RAD51 interface pulldowns (Figure 4).

Pulldown constructs were transformed into BL21 Rosetta (DE3) *E. coli* strains. Individual colonies were picked and grown overnight 25°C in 20 ml of Turbo broth expression media, supplemented with kanamycin and chloramphenicol. Resulting starter cultures were diluted

1:100 in 500 ml of Turbo Broth with kanamycin and chloramphenicol and grown to OD ~ 2.5. Cultures were cooled to room temperature before induction with 200 μM IPTG and expression at 15 °C for 16 hrs. Cultures were collected by centrifugation and resuspended in resuspension buffer (20 mM Tris pH8.0, 500 mM NaCl, and 20 mM Imidazole). Samples were either directly lysed or flash frozen with liquid N2. Lyses was performed by an emulsiflex C-5 (avestin) homogenizer, followed by centrifugation (50,000*g* for 1 hr). The protein complexes were pull-downed by gravity over 1 ml Ni-NTA column (Qiagen) in series with 0.5 ml Streptactin XT column (fig. S2B). After 3 column volume (CV) washes with resuspension buffer, both columns were then eluted by gravity separately with resuspension buffer +200 mM imidazole (Ni-NTA) or resuspension buffer + 100 mM biotin (Streptactin). Peak elution fractions were further purified by gravity over 1 ml amylose resin and eluted with resuspension buffer + 20 mM maltose, collected in 0.5 CV fractions. All elution fractions were confirmed by SDS-PAGE with Coomassie staining.

**Electrophoresis mobility shift assay (EMSA)**

DNA oligos were dissolved in 20 mM Tris pH 8.0 and 150 mM KCl to a final concentration of 200 μM if subsequent annealing was intended, or 100 μM if intended to remain single stranded. A ~50 ml 1% agarose gel in 0.5X TBE buffer (Life Technologies) was equilibrated at 4°C and pre-run at 30V for 1 hour. Isolated Rad51 Dimer and TR2i-RAD51 Dimer samples were each incubated in 5:1 ratio (RAD51:DNA), with 10 μM of various fluorescein-labeled DNA oligos in 10 μl total volumes (20mM Tris pH 8.0, 150 mM KCl) at 4°C for 15 minutes under dark. Subsequently, 1 μl of loading buffer (20 mM Tris pH 8.0, 40% glycerol) was added to each sample before being loaded on the agarose gel, and subsequently run at 30V at 4°C for 2.5 hours (covered to minimize light exposure). DNA electromobility was visualized by exposuring to UV radiation.

**Cell Culture and Transfections**

HeLa Kyoto WT, BRCA2 mutant and RAD51 mutant cells were cultured in Dulbecco's modified Eagle's medium (DMEM, Gibco) supplemented with 10% FBS, 100 U/ml penicillin and streptomycin and maintained at 37°C with 5% $CO_2$. JetPRIME® transfection reagent (#101000046) was used for all transfections according to the manufacturer's instruction.

**Generation of BRCA2 and RAD51 mutated cell lines**

sgRNAs were designed and selected for higher efficiency and reduced off-targets using sgRNA designer tool (https://portals.broadinstitute.org/gpp/public/analysis-tools/sgrna-design). The oligonucleotides used for targeted cleavage are: RAD51 M210L

(Top- 5' CACCGTTATCAAGCATCAGCCATGA 3' and

Bottom- 5' AAACTCATGGCTGATGCTTGATAAC 3'),

RAD51 M210I (Top- 5' CACCGTTATCAAGCATCAGCCATGA 3' and Bottom- 5' AAACTCATGGCTGATGCTTGATAAC 3'),

and BRCA2 P3292L (Top- 5' CACCGCATTTGTACATTTGTTTCTC 3' and Bottom- 5' AAACGAGAAACAAATGTACAAATGC 3').

Each oligonucleotide pair was phosphorylated and annealed at 37°C for 30 min; followed by 95°C for 5 min; ramp down to 25°C at 5°C/min in a thermocycler (T100, Bio-Rad). The phosphorylated and annealed oligos are diluted at 1:200 and cloned into *BbsI* sites of pSpCas9(BB)-2A-GFP (PX458) plasmid (Addgene Plasmid #48138). The cloned vectors were pretreated with PlasmidSafe exonuclease and transformed into ice-cold chemically competent Stbl3 cells according to manufacturer's protocol. The transformed cells were spread on LB plate containing ampicillin (100 µg/mL) and incubated overnight at 37°C. Positive colonies were picked and screened for sgRNA expression plasmid by sequencing (1$^{st}$ BASE) (fig. S3).

Single-stranded oligo DNA oligonucleotides (ssODNs) containing the desired point mutation along with a silent mutation (to incorporate a diagnostic restriction site) were synthesized by Integrated DNA Technologies (IDT) as custom-made 180-bp long oligos and used as HDR templates (See Table S2). 0.5 µg gRNA expression plasmid along with 10 µM HDR template were transfected into HeLa Kyoto cells. After 24 h, cells were sorted for GFP expression using the BD FACS Aria cell sorter into 96-well plate. ~2 weeks later, surviving single cell clones were expanded and screened by PCR amplifying targeted regions followed by diagnostic restriction digestion of the PCR product. Positive single cell colonies were further analyzed by sequencing of PCR products and cloning of PCR product into pCR Blunt vector (Invitrogen) followed by DNA sequencing. Cell line used in this studies are listed in Table S3.

**mClover Lamin A HDR assay**

Cells grown on coverslips in six-well plates were transfected with sgRNA plasmid targeting Lamin A (pUC CBASpCas9.EF1a-BFP.sgLMNA, Addgene Plasmid #98971) and donor plasmid (pCAGGS Donor mClover-LMNA, Addgene Plasmid #98970). Three days later, cells were fixed with 4% PFA (Sigma #252549) for 10 min and permeabilized with TBS Triton buffer for 5 min before mounting with DAPI containing medium. Images acquired with LSM 880 microscope were analyzed with ImageJ. HDR positive cells were defined as cells with mean mClover nuclear intensity over a threshold set for each experiment

**Olaparib Sensitivity Assay**

HeLa cells were seeded in 96-well plates and following overnight incubation, cells were treated with PARP1 inhibitor olaparib (Selleck Chemical #S1060) for 72 h at the indicated concentrations. Cell growth was then determined using CellTiter 96® AQueous One Solution as per the manufacturer’s instructions. Briefly, cell titer reagent (MTS with phenazine ethosulfate) is directly added to each well and incubated for 2 h at 37°C. Absorbance was recorded at 490nm in a 96 well plate reader (Tecan Spark® Multimode Microplate Reader).

**DNA Fiber Assay**

Briefly, HeLa cells were labelled with IdU (25 μM) (Sigma #I7125) for 20 min followed by 4 mM hydroxyurea (HU) (Sigma #H8627) treatment for 5 h. Cells were then harvested, resuspended in ice-cold PBS and spotted onto glass slides and lysed (200 mM Tris-HCl, pH 7.4, 50 mM EDTA, 0.5% SDS). DNA fibers were spread by tilting slides at 15-45° angle, followed by drying and fixing in 3:1 methanol/acetic acid for 10 min. The resulting DNA spreads were air-dried at room temperature overnight and denatured in 2.5 M HCl for 1 h followed by washing in ice-cold PBS three times. Slides were then blocked in 1.5% blocking buffer (Roche #11096176001) in 1X PBS/0.05% T20 for 30 min at 37°C. IdU tracks was detected using mouse anti-BrdU (BD #347580, clone B44; 1:5; 45 min at RT), followed by sequential labelling with secondary antibody (Rabbit anti-mouse AF594; 1:50; 20 min at RT) and tertiary antibody (Goat anti-rabbit AF594; 1:50; 20 min at RT). Single-stranded DNA was detected using mouse anti-ssDNA (MAB3034, Merck Millipore; 1:25; 45 min at RT) followed by sequential labelling with secondary antibody (rabbit anti-mouse AF488; 1:100; 20 min at RT) and tertiary antibody (donkey anti-rabbit AF488; 1:100; 20 min at RT). Each antibody incubation was followed by three PBS washes. Fibers were visualized by mounting slides in 90% glycerol in 1X PBS and imaged using LSM880 confocal microscope. Tract lengths were analyzed by ImageJ software.

**Direct stochastic optical reconstruction microscopy (dSTORM)**

Briefly, cells were seeded in 8 well chamber (Lab-Tek II Chambered Cover Glass W/Cover #1.5 Borosilicate #155409) and were treated with Neocarzinostatin (NCS) for 1h followed by recovery for 3 h and 5 h respectively. Cells were washed once with ice cold solution PBS solution followed by fixation in 2% formaldehyde (Sigma #252549) at room temperature (RT) for 20 min, then washed three times in PBS. Cells were then incubated with 50 mM $NH_4Cl$ (Sigma # A9434) for 30 min at RT. Next, cells were permeabilized with 0.2% Triton X-100 (Sigma #93443) for 10 min at RT. Nonspecific protein-protein interactions were blocked with 2% bovine serum albumin, 0.1% Triton X-100 and 0.05% Tween (Sigma #11332465001) in PBS for 1h. Primary antibodies (Rabbit anti-RAD51 monoclonal antibody (Abcam, ab133534, 1:2000) and Mouse anti-RPA32 monoclonal antibody (Abcam, clone 9H8, ab2175, 1:250) were incubated in blocking solution at 4°C overnight in a humidified chamber. RAD51 was fluorescently stained with goat anti-rabbit (Invitrogen, A-21246) Fab2 secondary antibody fragment conjugated to Alexa Fluor 647 for 1 h and RPA was detected with goat anti-mouse (Sigma, SAB4600401) CF568 conjugated Fab2 secondary antibody fragment. The fiducial marker, TetraSpeckTM beads (Invitrogen #T7279) were added to the fixed cells at 1:100 dilution and washed 3 times with PBS after 15 min incubation. The samples were kept at 4 °C in PBS until STORM imaging was performed.

STORM imaging was performed as previously described [59]. Briefly, imaging buffer containing 1% β-mercaptoethanol, 10% glucose, 0.5 mg/mL glucose oxidase and 40 μg/mL catalase were prepared and added freshly to the sample prior to imaging. STORM imaging was performed at room temperature on an inverted TIRF microscope (N-STORM, Nikon) with an Apochromat 100x/1.49 NA oil immersion objective lens. Images were acquired under highly inclined illumination mode 3 on a piezo stage coupled with Nikon Perfect Focus System for minimal z-focus drift. The STORM image acquisition was done sequentially for 647 nm channel followed by 561 nm channel. For 647 nm channel, we acquired 40,000 frames with a 20 ms exposure time at a power density of ~3 kW/cm2. For 561 nm channel, we acquired 40,000 frames with a 30 ms exposure time at a power density of ~6 kW/cm2. The fiducial markers for drift correction were imaged for 3 frames every 1000 frames. Images were acquired using an EMCCD camera (iXon Ultra DU897, Andor). The resolution achievable with our STORM imaging system is ~25 nm as previously measured 2.

Single molecule localization analysis was performed using custom-written codes in ImageJ and Matlab. Image stacks were first subjected to a wavelet-based denoising algorithm4. For each frame, individual spots were isolated by segmentation with a watershed algorithm and the coordinates of their center were determined using a radial gradient algorithm5, which is as accurate as a gaussian fit but much faster. The fiducial marker trajectories were extracted with a custom-written script to correct for lateral drift of the sample as well as to register the two channels. Single-molecule localizations with at least 50 photon counts were selected for further processing. Isolating individual RAD51 filaments was achieved using a combination of Delaunay triangulation and Voronoi diagram. Briefly, we first set a maximal distance between two nearest points via Delaunay triangulation. The background detection is removed by ignoring all segments whose length is beyond this threshold. This step alone is sufficient to isolate RAD51 filaments. To improve the delineation of the filaments, we set a maximal area via the Voronoi diagram to remove outlier points. Finally, the RAD51 filament length was obtained by determining the longest path among all the shortest paths connecting any two points within the filament.

**In situ protein interactions with nascent DNA replication forks (SIRF) assay**

SIRF assays were conducted as previously described [60,61] Briefly, cells were pulse treated with 125 µM EdU for 8 min, followed by 2 mM hydroxyurea for the indicated time. Cells were fixed, permeabilized and click-iT reaction was performed using biotin azide and AlexaFluor 488 azide according to manufacturers' instructions. Subsequently, proximity ligation assay (PLA) was performed using antibodies against the indicated RAD51/BRCA2 variant and biotinylated EdU. SIRF signals were imaged using the Nikon Eclipse Ti inverted microspore and analyzed using Nikon NIS-elements software. Antibodies used in SIRF are rabbit anti-biotin (Cell Signaling Technology, D5A7), and mouse anti-RAD51 (Abcam, ab213).

**Statistics and reproducibility**

Statistical analysis was conducted using GraphPad Prism software (v10). Statistical tests used are listed in the relevant figure legends. P-value summary for all statistical tests used in this study is: ns, P-value > 0.05; * P-value ≤ 0.05; ** P value ≤ 0.01; *** P-value ≤ 0.001; **** P-value ≤ 0.0001. Experiments were done independently at least three times.

**Data and materials availability**

The crystal structure of TR2i-RAD51 dimer complex has been deposited to the Protein Data Bank (PDB). PDB ID: 8UVW.


69. Otwinowski, Z., and Minor, W. (1997). Processing of X-ray diffraction data collected in oscillation mode. Methods Enzymol *276*, 307-326. 10.1016/S0076-6879(97)76066-X.

70. Agirre, J., Atanasova, M., Bagdonas, H., Ballard, C.B., Basle, A., Beilsten-Edmands, J., Borges, R.J., Brown, D.G., Burgos-Marmol, J.J., Berrisford, J.M., et al. (2023). The CCP4 suite: integrative software for macromolecular crystallography. Acta Crystallogr D Struct Biol *79*, 449-461. 10.1107/S2059798323003595.

71. Liebschner, D., Afonine, P.V., Baker, M.L., Bunkoczi, G., Chen, V.B., Croll, T.I., Hintze, B., Hung, L.W., Jain, S., McCoy, A.J., et al. (2019). Macromolecular structure determination using X-rays, neutrons and electrons: recent developments in Phenix. Acta Crystallogr D Struct Biol *75*, 861-877. 10.1107/S2059798319011471.

72. Emsley, P., and Cowtan, K. (2004). Coot: model-building tools for molecular graphics. Acta Crystallogr D Biol Crystallogr *60*, 2126-2132. 10.1107/S0907444904019158.
